# Finding genetic variants in plants without complete genomes

**DOI:** 10.1101/818096

**Authors:** Yoav Voichek, Detlef Weigel

## Abstract

Structural variants and presence/absence polymorphisms are common in plant genomes, yet they are routinely overlooked in genome-wide association studies (GWAS). Here, we expand the genetic variants detected in GWAS to include major deletions, insertions, and rearrangements. We first use raw sequencing data directly to derive short sequences, *k*-mers, that mark a broad range of polymorphisms independently of a reference genome. We then link *k*-mers associated with phenotypes to specific genomic regions. Using this approach, we re-analyzed 2,000 traits measured in *Arabidopsis thaliana*, tomato, and maize populations. Associations identified with *k*-mers recapitulate those found with single-nucleotide polymorphisms (SNPs), however, with stronger statistical support. Moreover, we identified new associations with structural variants and with regions missing from reference genomes. Our results demonstrate the power of performing GWAS before linking sequence reads to specific genomic regions, which allow detection of a wider range of genetic variants responsible for phenotypic variation.

## Introduction

Elucidating the link between genotype and phenotype is central to biological research, both in basic research as well as in translational medicine and agriculture. Correlating genotypic and phenotypic variability in genome-wide association studies (GWAS) has become the tool of choice for systematic identification of candidate loci in the genome that are causal for phenotypic differences. In plants, many species-centered projects are genotyping collections of individuals, for which different phenotypes can then be measured and analyzed. These include hundreds or thousands of strains from *Arabidopsis thaliana*, rice, maize, tomato, sunflower, and several other species (1001 Genomes Consortium, 2016; Bukowski et al., 2018; Hübner et al., 2018; Tieman et al., 2017; Wang et al., 2018).

A difficulty of working with plant genomes is that they are highly repetitive and feature excessive structural variation between members of the same species, mostly attributed to their active transposons (Bennetzen, 2000). For example, in the well-studied species *Arabidopsis thaliana*, natural accessions are missing 15% of the reference genome, indicating a similar fraction would be absent from the reference, but present in other accessions (1001 Genomes Consortium, 2016). Moreover, although *A. thaliana* has a small (140 Mb) and not very repetitive genome compared to many other plants, SNPs may be assigned to incorrect positions due to sequence similarity shared between unlinked loci (Long et al., 2013). The picture is even more complicated in other plant species, such as maize. The maize 2.3 Gb genome is highly repetitive, with transposons often inserted into other transposons, and 50%-60% of short read sequences can not be mapped uniquely to it, making the accurate identification of variants in the population a formidable challenge (Bukowski et al., 2018; Schnable et al., 2009). Furthermore, about 30% of low-copy genes present in the entire population are not found in the reference (Gore et al., 2009; Springer et al., 2018; Sun et al., 2018). Presence of large structural variants are ubiquitous all over the plant kingdom, and there are many examples for their effects on phenotypes (Saxena et al., 2014). The importance of structural variants in driving phenotypic variation has been appreciated from the early days of maize genetics (McClintock, 1950), though searching for them systematically is still an unsolved problem.

Correlating phenotypic and genotypic variation in GWAS is critically dependent on the ability to call individual genotypes. While short sequencing reads aligned to a reference genome can identify variants smaller than read length, such as SNPs and short indels, this approach is much less effective for larger structural variants. Moreover, variants such as SNPs can be in regions missing from the reference genome, which is frequently the case in plants. Organellar genomes are a special case, being left out of GWAS systematically although their genetic variation was shown to have strong phenotypic effects (Davila et al., 2011; Joseph et al., 2013). Although not regularly used, short read sequencing can provide, in principle, information for many more variants in their source genomes than only SNPs and short indels (Iqbal et al., 2012).

While variants are typically discovered with short reads by mapping them to a target reference genome, one can also directly compare common subsequences among samples (Zielezinski et al., 2019). Such a direct approach is intuitively most powerful when the reference genome assembly is poor, or even non-existent. Because short reads result from random shearing of genomic DNA, and because they contain sequencing errors, comparing short reads between two samples directly is, however, not very effective. Instead, genetic variants in a population can be discovered by focusing on sequences of constant length *k* that are even shorter than typical short reads, termed *k*-mers. After *k*-mers have been extracted from all short reads, sets of *k*-mers present in different samples can be compared. Importantly, *k*-mers present in some samples, but missing from others, can identify a broad range of genetic variants. For example, two genomes differing in a SNP (Fig. 1A) will have *k k*-mers unique to each genome; this is true even if the SNP is found in a repeated region or a region not found in the reference genome. Structural variants, such as large deletions, inversions, translocations, transposable element (TE) insertion, etc. will also leave marks in the presence or absence of *k*-mers (Fig. 1A). Therefore, instead of defining genetic variants in a population relative to a reference genome, a *k*-mer presence/absence in raw sequencing data can be directly associated with phenotypes to enlarge the tagged genetic variants in GWAS (Lees et al., 2016).

**Figure 1.**
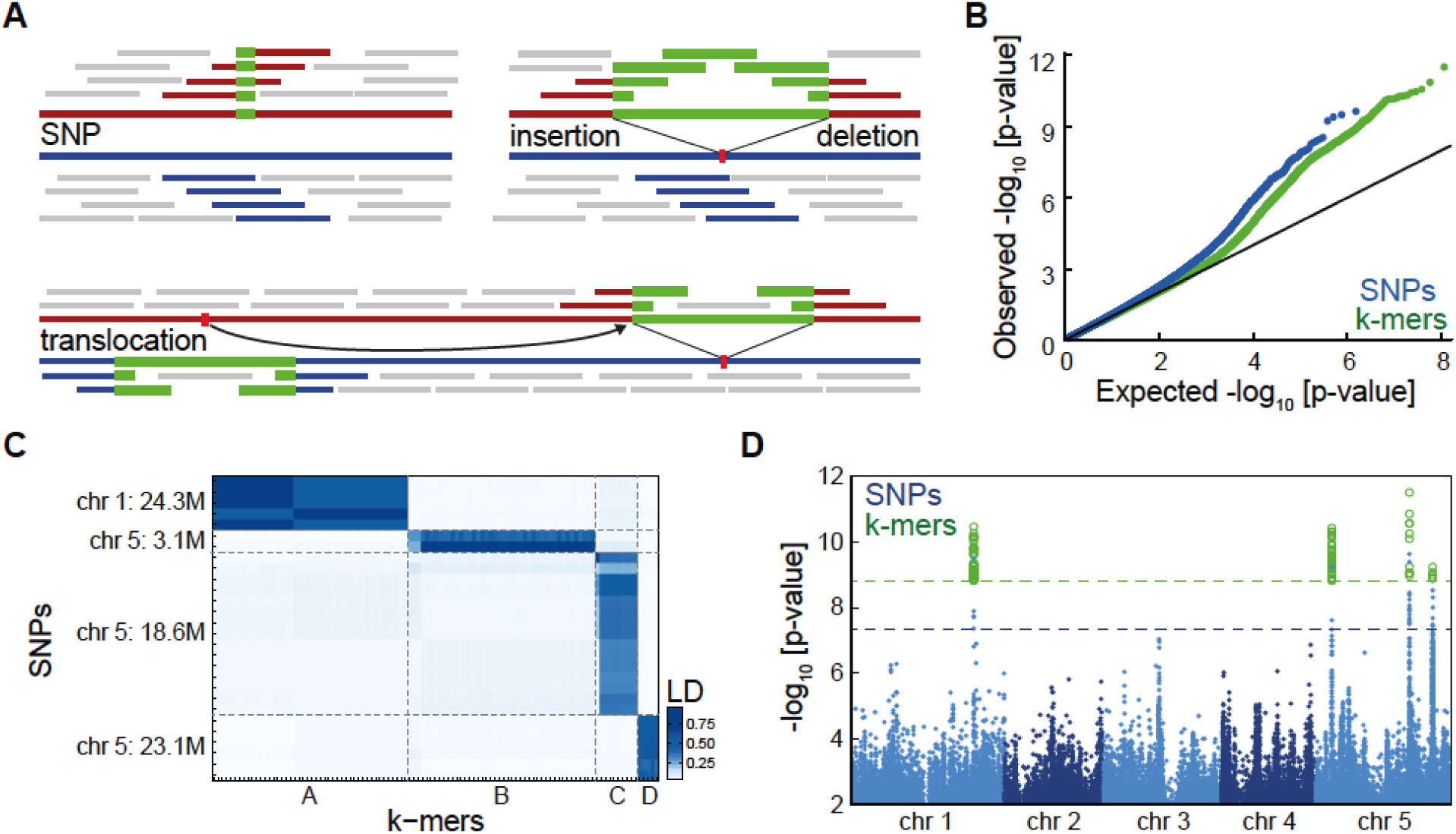
Flowering time associations in *A. thaliana*. **(A)** Presence and absence of *k*-mers marks a range of different genetic variants. Blue and red lines represent two individuals genomes, and short bars above/below mark in color the *k*-mers unique to each genome due to genomic differences or in grey ones shared between genomes. **(B)** P-values quantile-quantile plot of SNPs and *k*-mers associations with flowering time measured in 10°C. Deviation from the black line (y=x) indicate stronger associations than expected by chance. **(C)** LD (expressed as r^2^) between all SNPs and *k*-mers passing the p-value thresholds. Four highly linked families of variants were identified with both methods. For SNP-to-SNP and *k*-mer-to-*k*-mer LD, see Fig. S2B,C. **(D)** Manhattan plot showing p-values of all SNPs (blue) and of the subset of *k*-mers passing the p-value threshold (green) as a function of their genomic position. Dashed lines mark the p-value thresholds for SNPs (blue) and *k*-mers (green).

Reference-free GWAS based on *k*-mers has been used for mapping genetic variants in bacteria, where each strain contains only a fraction of the genes present in the pan-genome (Lees et al., 2016, 2017; Sheppard et al., 2013). This approach, not centered around one specific reference genome, can identify biochemical pathways associated with, for example, pathogenicity. This approach has also been applied in humans, where the number of unique *k*-mers is much higher than in bacterial strains, due to their larger genome (Rahman et al., 2018). However, this was restricted to case-control situations, and due to high computational load, population structure was corrected only for a subset of *k*-mers.

While *k*-mer based approaches are likely to be especially appropriate for plants, the large genome sizes, highly structured populations, and excessive genetic variation (Gordon et al., 2017; Minio et al., 2019; Sun et al., 2018) limit the application of previous *k*-mer methods to plants. A first attempt to nevertheless use *k*-mer based methods has recently been made in plants, but was limited to a small subset of the genome, and also accounting for population structure only for a small subset of all *k*-mers (Arora et al., 2019).

Here, we present an efficient method for *k*-mer-based GWAS and compare it directly to the conventional SNP-based approach on more than 2,000 phenotypes from three plant species with different genome and population characteristics - *A. thaliana*, maize and tomato. Most variants identified by SNPs can be detected with *k*-mers (and vice versa), but *k*-mers having stronger statistical support. For *k*-mer-only hits, we demonstrate how different strategies can be used to infer their genomic context, including large structural variants, sequences missing from the reference genome, and organeller variants. Lastly, we compute population structure directly from *k*-mers, enabling the analysis of species with poor quality or without a reference genome. In summary, we have inverted the conventional approach of building a genome, using it to find population variants, and only then associating variants with phenotypes. In contrast, we begin by associating sequencing reads with phenotypes, and only then infer the genomic context of these sequences. We posit that this change of order is especially effective in plant species, for which defining the full population-level genetic variation based on reference genomes remains highly challenging.

## Results

### Proof of concept: genetic variants for flowering of *A. thaliana*

As an initial proof of concept, we looked at the well-studied and well-understood trait in the model plant *A. thaliana*, flowering time. In *A. thaliana*, GWAS approaches have been used for almost 15 years (Aranzana et al., 2005), and 1,135 individuals, termed accessions, had their entire genomes resequenced several years ago (1001 Genomes Consortium, 2016). We used this genomic dataset to define the presence/absence patterns of 31 bp *k*-mers in these accessions (Fig. S1A). In order to minimize the effect of sequencing errors, for each DNA-Seq dataset we only considered *k*-mers appearing at least thrice. Out of a total 2.26 billion unique *k*-mers across the entire population, 439 million appeared in at least five accessions (Fig. S2A). These *k*-mers were not shared by all accessions, and we used the presence or absence of a *k*-mer as two alleles per variant to perform GWA with a linear mixed model (LMM) to account for population structure (Fig. S1B) (Zhou and Stephens, 2012). For comparison purposes, GWA was performed also with SNPs and short indels. In both cases statistically significant associations were detected (Fig. 1B).

To define a set of *k*-mers most likely to be associated with flowering time, we had to set a p-value threshold. A complication in defining such a threshold is that *k*-mers are often not independent, as a single genetic variant is typically tagged by several *k*-mers (Fig. 1A). For example, 180 million *k*-mers had a minor allele frequency above 5%, but these represented only 110 million unique presence/absence patterns across accessions. Thus, a Bonferroni correction based on the number of all tests would be inaccurate, as it would not accurately reflect the effective number of independent tests. To define a threshold that accounts for the dependencies between *k*-mers we therefore used permutation of the phenotype (Abney, 2015). This approach presents a computational challenge, as the full GWA analysis has to be run multiple times. To this end, we implemented a LMM-based GWA specifically optimized for the *k*-mer application (Fig. S1C) (Loh et al., 2015; Svishcheva et al., 2012).

We calculated the p-value thresholds for SNPs and *k*-mers, set to a 5% chance of getting one false-positive. The threshold for *k*-mers was more stringent than the one for SNPs (35-fold), but lower than the increase in tests number (140-fold), as expected due to the higher dependency between *k*-mers. Twenty-eight SNPs and 105 *k*-mers passed their corresponding thresholds. Using LD, we linked SNPs to *k*-mers directly without locating the *k*-mers genomic locations. Four distinct families of linked genetic variants were identified in both methods, with a clear one-to-one relationship between the four sets of SNPs and the four sets of *k*-mers (Fig. 1C, Fig. S2B,C). As expected, the *k*-mers aligned to the same genomic loci as the corresponding SNPs (Fig. 1D). For validation, we ran the analysis again with a *k*-mer length of 25 bp, obtaining a very similar result (Fig. S2D). Therefore, in this case, *k*-mer based GWAS identified the same genotype-phenotype associations as detected by SNPs.

### Comparison of SNP- and *k*-mer-based GWAS on 1,697 *A. thaliana* phenotypes

Flowering time is a very well studied trait, and it is unlikely that a new locus affecting it will be discovered by GWAS. To assess the potential of *k*-mer-based GWA to identify new associations, we set out to systematically compare it to the SNP-based method on a comprehensive set of traits. To this end, we collected 1,697 phenotypes from 104 *A. thaliana* studies (Table S1). This collection spans a representative sample of phenotypes regularly measured in plants (Fig. 2A). Eliminating phenotypes for which there are short read sequencing data from fewer than 40 accessions, we were left with 1,582 traits to which both methods could be applied. All parameters affecting GWA analysis, such as minor allele frequency or relatedness between individuals, were the same, to obtain the most meaningful comparison. Moreover, as *A. thaliana* is a selfer, SNPs are homozygous, and their state is therefore comparable to the binary *k*-mer presence/absence.

**Figure 2.**
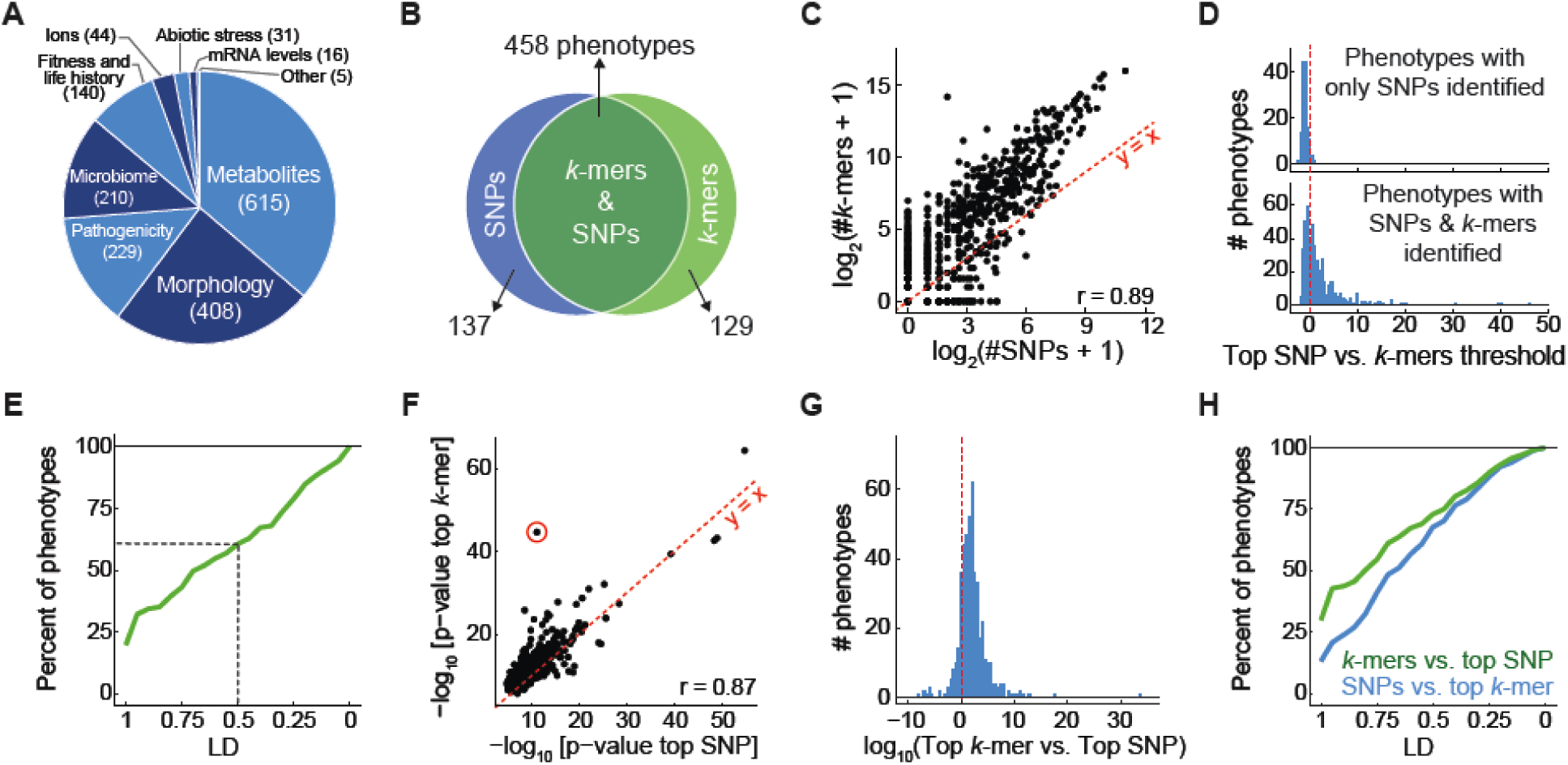
Comparison of SNP- and *k*-mer-based GWAS on 1,697 *A. thaliana* phenotypes. **(A)** Assignment of 1,697 phenotypes to broad categories. **(B)** Overlap between phenotypes with SNP and *k*-mer hits. **(C)** Correlation of number of significantly associated *k*-mers vs. SNPs for all phenotypes. **(D)** Ratios (in log_10_) of top SNP p-value vs. the *k*-mers threshold for 137 phenotypes with only significant SNPs (top), and for 458 phenotypes with both significant SNPs and *k*-mers (bottom). **(E)** Fraction of phenotypes, from 137 phenotypes that had only significant SNP hits, for which a *k*-mer passing the SNP threshold could be found within different LD cutoffs. For a minimum of LD=0.5 (dashed lines), 61% of phenotypes had a linked *k*-mer that passed the SNP threshold. **(F)** Correlation of p-values of top *k*-mers and SNPs for all phenotypes (r=0.87). Red circle marks the strongest outlier (see Fig. 3A, B for details on this phenotype). **(G)** Ratio between top p-values (expressed as -log_10_) in the two methods, for the 458 phenotypes with both *k*-mer and SNP hits. **(H)** Fraction of all phenotypes for which a significant SNP could be found within different LD cutoffs of top *k*-mer (blue) and vice versa (green).

We first wanted to learn whether the two methods identified similar associations. Indeed, there was substantial overlap between the traits for which associations were found (Fig. 2B). Also, the number of identified *k*-mers and SNPs per phenotype were correlated (r=0.89), and as expected, more associated *k*-mers than SNPs were identified (Fig. 2C, Fig. S3A). For 137 phenotypes, only a significant SNP could be identified, due to the more stringent thresholds for *k*-mers, as the most significant SNPs in almost all of these phenotypes did not pass the *k*-mer threshold (Fig. 2D). Moreover, in most of these phenotypes, a *k*-mer passing the SNPs threshold was in high LD with the top SNP (Fig. 2E). Although the *k*-mer thresholds were more stringent than the SNPs thresholds (Fig. S3B), for 129 phenotypes only *k*-mers but no SNPs associations were identified. These cases were the best candidates for associations that cannot be captured with SNPs.

We next compared p-values of top SNPs to those of top *k*-mers; the two were correlated (r=0.87, Fig. 2F). Focusing on phenotypes for which both SNPs and *k*-mers were identified, the great majority, 86%, had stronger p-values for the top *k*-mer (Fig. 2G), a trend that had already been observed for flowering time (Fig. 1D). Lastly, we wanted to know how well top *k*-mers were tagged by significantly associated SNPs and vice versa. We quantified this with the LD (as in Fig. 1C) between the top SNP and the closest associated *k*-mer and the other way around. While SNPs tagged variants similar to top *k*-mers, associated *k*-mers were on average closer to top SNPs than associated SNPs to top *k*-mers (Fig. 2H). This was expected, as *k*-mers can represent SNPs but also capture other types of genetic variants.

### Specific case studies of *k*-mer superiority

For some phenotypes, *k*-mers were more strongly associated with a phenotype than the top SNP, although they represented the same variant (Fig. S4A). The goal of our study was, however, to identify cases where *k*-mers provided a conceptual improvement. First, we looked into the phenotype quantifying the fraction of dihydroxybenzoic acid (DHBA) xylosides among total DHBA glycosides (Li et al., 2014) (red circle in Fig. 2F). In this case, all significant *k*-mers mapped uniquely in the proximity of AT5G03490, encoding a UDP glycosyltransferase that was identified in the original study as causal (Fig. 3A, Fig. S4B). The source of the stronger *k*-mers associations could be traced back to two non-synonymous SNPs, 4 bp apart, in the coding region of AT5G03490. Due to their proximity, one *k*-mer can hold the state of both SNPs, and their combined information is more predictive of the phenotype than each SNP on its own (Fig. 3B). This interaction between closely linked SNPs was not one of the types of genetic variants we had anticipated for *k*-mers.

**Figure 3.**
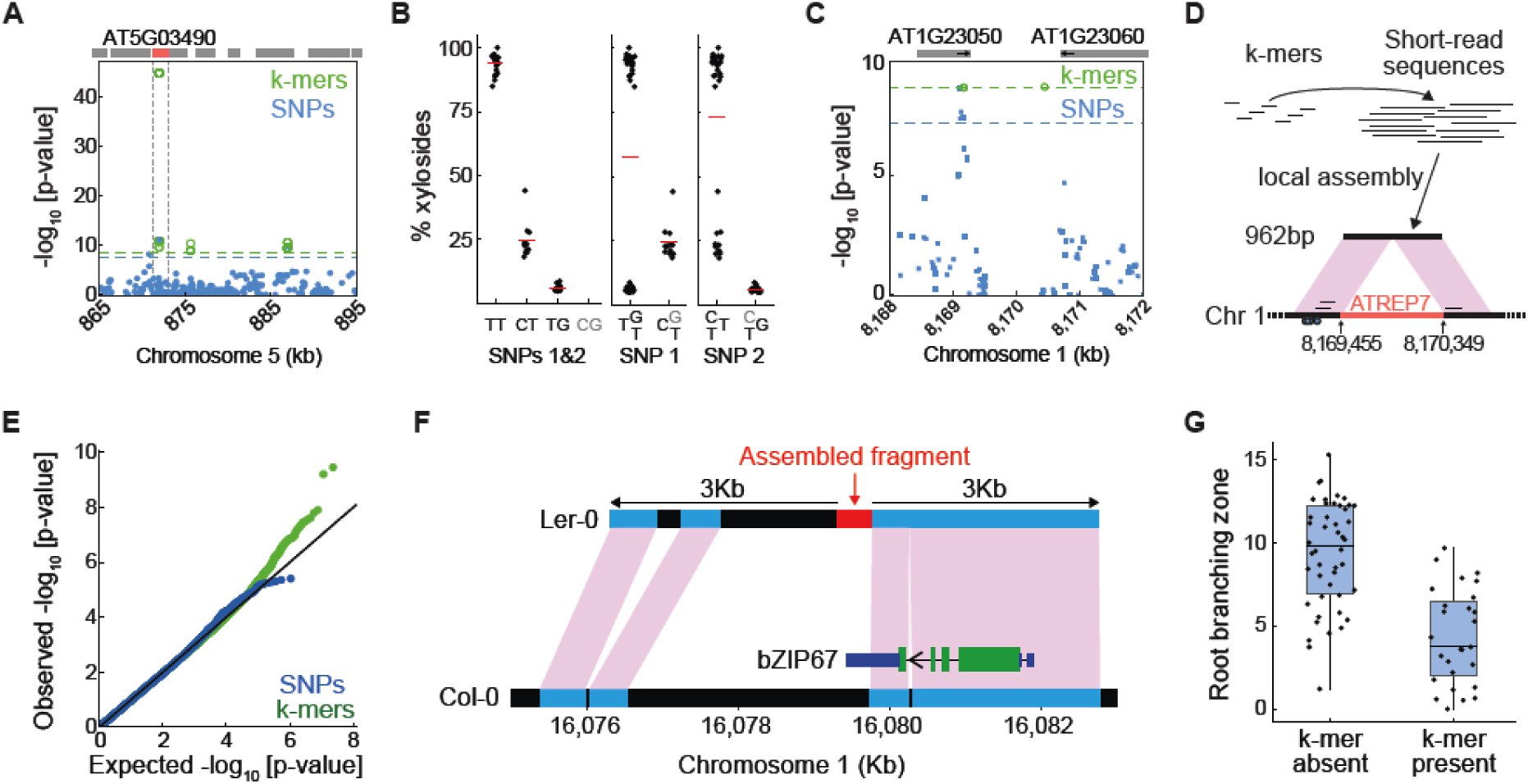
Specific cases in which *k*-mers are superior to SNPs. **(A)** Associations with xyloside fraction in a region of chromosome 5. Grey boxes indicate genes with AT5G03490 marked in red. **(B)** Xyloside fraction grouped by states at two SNPs (SNP1, 872,003 bp; SNP 2, 872,007 bp). One of the four possible states (“CG”) does not exist, indicated in grey in left most plot, which shows grouping based on both sites, as is possible with *k*-mers. Middle and right most plot show groupings based on only one of the two sites. **(C)** Associations with seedling growth inhibition in the presence of flg22 near 8.17 Mb of chromosome 1. Absence of SNPs in the central 1 kb region is likely due to the presence of a TE to which short reads cannot be unambiguously mapped. Gene orientations indicated with short black arrows. **(D)** Assembly of reads identified with the seven unmappable *k*-mers resulted in a 962bp fragment. This fragment lacks the central 892 bp region in the reference genome encoding an ATREP7 helitron TE. Small circles on bottom represent significant flanking SNPs, and short black bars above represent the three mappable significant *k*-mers. **(E)** P-values quantile-quantile plot of associations with germination time in darkness and low nutrients. Only *k*-mers show stronger-than-expected associations. **(F)** Assembled reads (red bar) containing significant *k*-mers from GWA of germination time match a region on chromosome 3 of Ler-0. Regions in addition to the red fragment that cannot be aligned to the Col-0 reference genome are indicated in black. The 3’ UTR of the gene encoding bZIP67 is indicated in dark blue. The extent of the bZIP67 3’ UTR in Ler-0 is not known. Green indicates coding sequences. **(G)** Root branching zone length in millimeters in accessions that have the significant *k*-mer identified for this trait and accessions that do not have this *k*-mer.

Our next case study involves inhibition of seedling growth in the presence of a specific flg22 variant (Vetter et al., 2016), a phenotype for which we could map to the reference genome only three of the 10 significant *k*-mers; the three mappable *k*-mers were all located in the proximity of significant SNPs in AT1G23050 (Fig. 3C, Fig. S4C). To identify the genomic source of the remaining seven *k*-mers, we retrieved the short reads containing the *k*-mers from all relevant accessions and assembled them into a single 962 bp fragment. This fragment mapped to two genomic regions 892 bp apart, close to the three mapped *k*-mers (Fig. 3D). The junction sequence connecting the two regions could only be identified in accessions with the seven significant *k*-mers. We hypothesized that the 892 bp intervening fragment corresponds to a transposable element (TE), and a search of the Repbase database indeed identified similarity to helitron TE (Bao et al., 2015). Thus, the *k*-mers in this case marked an association with a structural variant, the presence or absence of a ∼900 bp TE. While in this case the *k*-mer method did not identify a new locus, it more clearly revealed what is the likely genetic cause of variation in flg22 sensitivity.

In the first two examples, hits with both *k*-mers and SNPs had been identified. Next, we looked for phenotypes for which we had only identified significant *k*-mers. One of these was germination in darkness and under low nutrient supply (Morrison and Linder, 2014). In this case, 11 *k*-mers but no significant SNPs had been found (Fig. 3E, Fig. S4D-E). However, neither the 11 *k*-mers nor the short reads they originated from could be mapped to the reference genome. The reads assembled into a 458 bp fragment. A database search revealed a hit on chromosome 3 of Ler-0, a non-reference accession of *A. thaliana* with a high-quality genome assembly (Zapata et al., 2016). The flanking sequences were syntenic with region on chromosome 3 of the *A. thaliana* reference genome, with a 2 kb structural variant that included the 458 bp fragment we had assembled based on our *k*-mer hits (Fig. 3F). This variant affected the 3’ untranslated region (UTR) of the bZIP67 transcription factor gene. bZIP67 acts downstream of LEC1 and upstream of DOG1, two master regulators of seed development (Bryant et al., 2019). Accumulation of bZIP67 protein but not *bZIP67* mRNA is affected by cold and thus likely mediates environmental regulation of germination (Bryant et al., 2019). Structural variations in the 3’ UTR is consistent with translational regulation of bZIP67 being important. The bZIP67/germination case study demonstrates directly the ability of our *k*-mer method to reveal associations with structural variants that are not tagged by SNPs.

As a final case, we focused on the variation in the length of the root branching zone (Ristova et al., 2018). While no significant SNPs could be identified, a single *k*-mer passed the significance threshold (Fig. 3G, Fig. S4F). The *k*-mer and the reads containing it mapped to the chloroplast genome. When we lowered the threshold for the familywise error-rate from 5% to 10%, a second *k*-mer was identified, which also mapped to the chloroplast genome. Genetic variation in organelle genomes has been shown to affect phenotypic variation (Joseph et al., 2013), but they are often left out from GWA studies.

### Comparison of SNP- and *k*-mer-based GWAS in maize

While the results with *A. thaliana* were encouraging, its genome size and repeat content is not representative of many other flowering plants. We therefore wanted to evaluate our approach on larger, more complex genomes. This criterion is met by maize, with a reference genome of 2.3 Gb, ∼85% of which consists of TEs and other repeats (Schnable et al., 2009). Moreover, individual maize genomes are highly divergent, with ∼10% of genes being non-syntenic and many genes found in different accessions are missing from the reference genome (Gore et al., 2009; Springer et al., 2018; Sun et al., 2018).

We set out to apply our *k*-mer-based GWAS approach to a set of 150 maize inbred lines with short read sequence coverage of at least 6x (Bukowski et al., 2018). There were 7.3 billion unique *k*-mers in the population, of which 2.3 billion were present in at least five accessions, which were used for GWAS (Fig. S5A). As in *A. thaliana*, we sought to compare the *k*-mer- and SNP-based approaches. To this end, we applied both methods to 252 field measurements, mostly of morphological traits (Zhao et al., 2006). For 89 traits, significant associations were identified by at least one of the methods, and for 37 by both (Fig. 4A). As in *A. thaliana*, the number of statistically significant variants as well as top associations between both methods were well correlated (Fig. 4B,C). Top *k*-mers had lower p-values than the top SNPs (Fig. S5D), and the *k*-mer method detected associations not found by SNPs.

**Figure 4.**
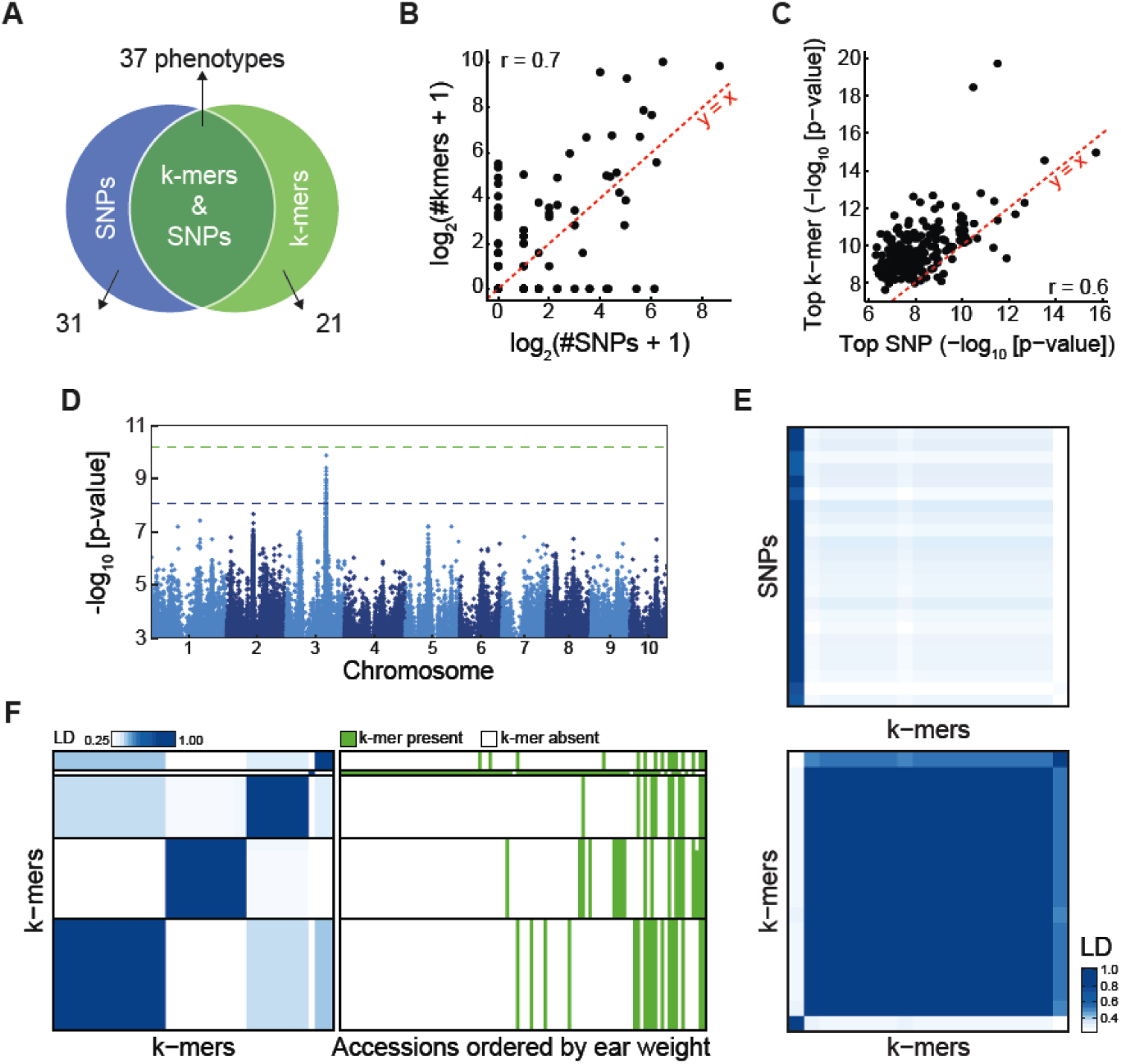
Comparison of SNP- and *k*-mer-based GWAS in maize. **(A)** Overlap between phenotypes with SNP and *k*-mer hits. See also Fig. S5B,C. **(B)** Correlation of number of significantly associated *k*-mers vs. SNPs for all phenotypes. See also Fig. S5E. **(C)** Correlation of p-values of top *k*-mers and SNPs for all phenotypes. **(D)** Manhattan plot of SNP associations with days to tassel (environment 06FL1). **(E)** LD between 23 significant SNPs and 18 *k*-mers (top) or *k*-mers to *k*-mers (bottom) for days to tassel. Order of *k*-mers is the same in both heatmaps. **(F)** LD between 45 *k*-mers associated with ear weight (environment 07A, left), and *k*-mer’s presence/absence patterns in different accessions ordered by their ear weight (right).

To discern the added benefit of the *k*-mer-based approach, we compared SNPs and *k*-mers using LD, without attempting to locate *k*-mers in the genome. We used this comparison approach as SNPs were originally placed on the genomic map using external information in addition to short read mapping, due to the large proportion of short reads that do not map to unique places in the reference genome (Bukowski et al., 2018). We found several cases where a *k*-mer marked a common allele in the population with strong effect on a phenotype, but the allele could not be identified with the SNP dataset. For example, for days to tassel there was one clear SNP hit that was also tagged by *k*-mers (Fig. 4D,E), but a second genetic variant was only identified with *k*-mers. Another example is ear weight for which no SNPs passed the significance threshold (Fig. S5F), but several unlinked variants were identified with *k*-mers (Fig. 4F). Thus, new alleles with high predictive power for maize traits can be revealed using *k*-mers.

A major challenge in identifying causal variants in maize is the high fraction of short reads that do not map uniquely to the genome. In the maize HapMap project, additional information had to be used to find the genomic position of SNPs, including population LD and genetic map position (Bukowski et al., 2018). The same difficulty of unique mappings also undermined the ability to identify the genomic source of *k*-mers associated with specific traits. For example, we tried to locate the genomic position of the *k*-mer corresponding to the SNP associated with days to tassel in chromosome 3 (Fig. 4D). The vast majority of short reads from which the *k*-mer originated, 99%, could not be mapped uniquely to the reference genome. However, when we assembled all these reads into a 924 bp contig, this fragment could now be uniquely placed in the genome, to the same place as the identified SNPs. Thus, as we were only interested in finding the genomic position after we already had an association in hand, we could use the richness of combining reads from many accessions to more precisely locate their origin without the use of additional genetic information, as had to be used for the SNPs.

### Comparison of SNP- and *k*-mer-based GWAS in tomato

Tomato has a 900 Mb genome, which is intermediate between *A. thaliana* and maize, but it presents its own challenges, as modern tomatoes show a complex history of recent introgressions from wild relatives (Lin et al., 2014; Tomato Genome Consortium, 2012). Of 3.2 billion unique *k*-mers in all 246 used accessions, 981 million were found in at least five accessions (Fig. S6A). We compared *k*-mer- and SNP-based GWAS on 96 metabolites measurements from two previous studies (Tieman et al., 2017; Zhu et al., 2018). For most metabolites, an association was identified by at least one method, with three metabolites having only SNP hits and 13 only *k*-mer hits (Fig. 5A). Similar to *A. thaliana* and maize, the number of identified variants as well as top p-values were correlated between methods (Fig. 5B,C). Top *k*-mers associations were also stronger than top SNPs (Fig. S5D), but even more so than in *A. thaliana* or maize, with an average difference of 10^4.4^, suggesting that in tomato the benefits of *k*-mer-based GWAS are also larger.

**Figure 5.**
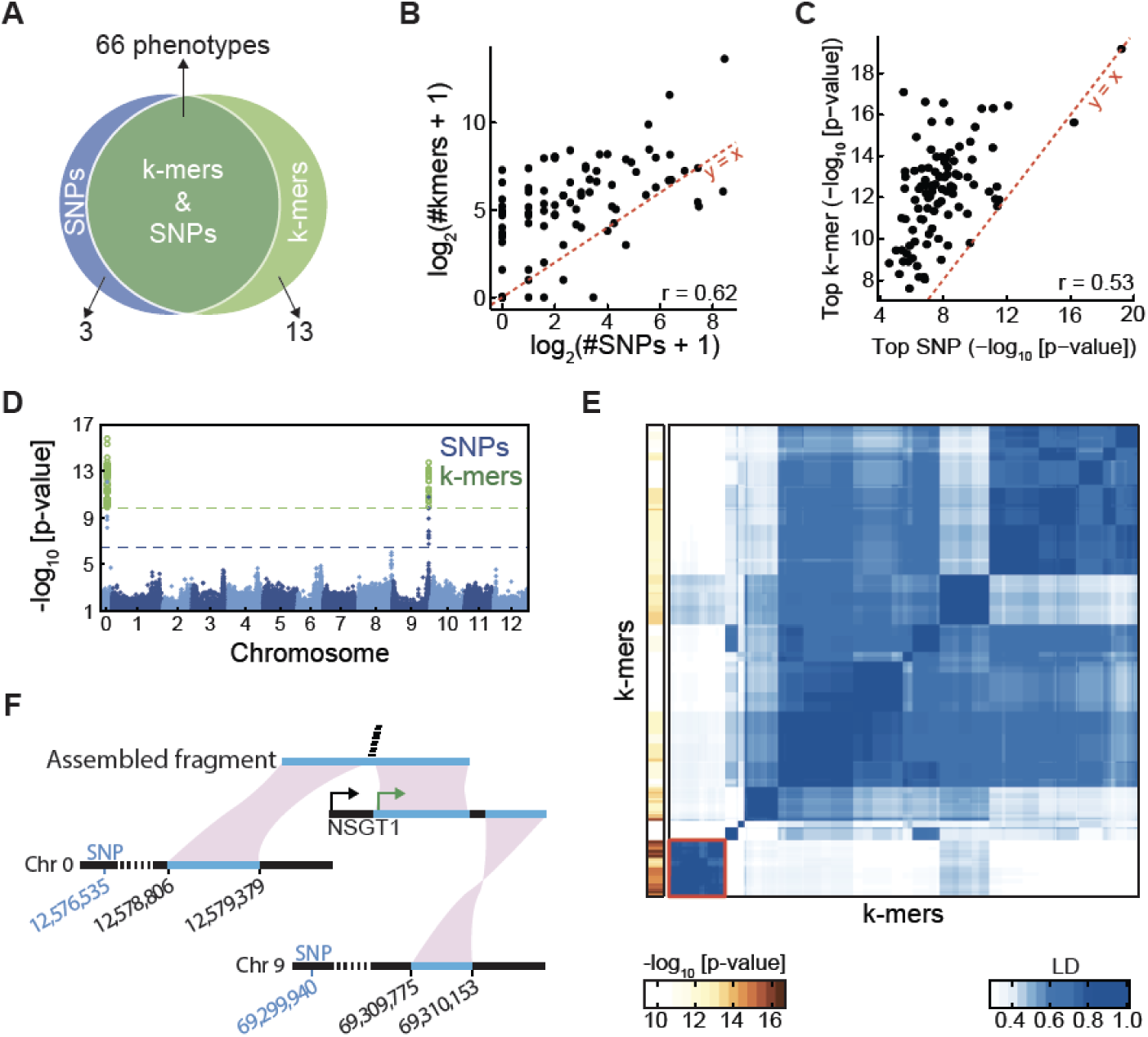
Comparison of SNP- and *k*-mer-based GWAS in tomato. **(A)** Overlap between phenotypes with SNP and *k*-mer hits. See also Fig. S5B,C. **(B)** Correlation of number of significantly associated *k*-mers vs. SNPs for all phenotypes. See also Fig. S5E. **(C)** Correlation of p-values of top *k*-mers and SNPs for all phenotypes. **(D)** Manhattan plot of SNPs and *k*-mers associations with guaiacol concentration. **(E)** LD among 293 *k*-mers associated with guaiacol concentration (right), and the p-value of each *k*-mer (left). Red square on bottom left indicates the 35 *k*-mers with strongest p-values and no mappings to the reference genome. **(F)** The first part of a fragment assembled from the 35 unmapped *k*-mers (E) mapped to chromosome 0 and the second part to the unanchored complete *NSGT1* gene. Only the 3’ end of *NSGT1* maps to the reference genome, to chromosome 9. The green and black arrows marks the start of the *NSGT1* ORF in the R104 “smoky” line and “non-smoky” lines, respectively (Tikunov et al., 2013). Two SNPs are indicated, which are the significant SNPs closest to the two regions of the reference genome.

We next looked, as a case-study, at measurements of guaiacol, which results in a strong off-flavor and is therefore not desirable (Tieman et al., 2017). SNPs in two genomic loci were associated with it (Fig. 5D), one in chromosome 9 and the other in what is called “chromosome 0”, which corresponds to the concatenation of all sequence scaffolds that could not be assigned to one of the 12 nuclear chromosomes. From the 293 significant *k*-mers, 180 could be mapped uniquely to the genome, all close to significant SNPs. Among the remaining *k*-mers, of particular interest was a group of 35 *k*-mers in very high LD that had the lowest p-values, but could not be mapped to the reference genome (Fig. 5E). Assembly of the reads containing these *k*-mers resulted in a 1,172 bp fragment, of which the first 574 bp could be aligned near significant SNPs in chromosome 0 (Fig. 5F). The remainder of this fragment could not be placed in the reference genome, but there was a database match to the *NON-SMOKY GLYCOSYLTRANSFERASE 1* (*NSGT1*) gene (Tikunov et al., 2013). The 35 significant *k*-mers covered the junction between these two mappable regions. Most of the *NSGT1* coding sequence is absent from the reference genome, but present in other accessions. *NSGT1* had been originally isolated as the causal gene for natural variation in guaiacol levels (Tikunov et al., 2013). Since *NSGT1* can be anchored to chromosome 9 near the identified SNPs (Fig. 5F), the significant SNPs identified in chromosomes 0 and 9 apparently represent the same region, connected by the fragment we assembled from our set of 35 significant *k*-mers. Thus, we identified an association outside the reference genome, and linked the SNPs in chromosome 0 to chromosome 9.

### Calculation of relatedness between individuals based on *k*-mers

We have shown that we can assemble short fragments from *k*-mer-containing short reads and find hits not only in the reference genome, but also in other published sequences. This opens the possibility to apply our *k*-mer-based GWAS method to species without a high-quality reference genome. Draft genomes with contigs that include typically multiple genes can be relatively easily and cheaply generated using short read technology (Sohn and Nam, 2018). The major question with such an approach is then how one would correct for population structure in the GWAS step.

So far, we had relied on SNP kinship information. If one were to extend our method to species without high-quality reference genomes one would ideally be able to learn kinship directly from *k*-mers, thus obviating the need to map reads to a reference genome for SNP calling. With this goal in mind, we estimated relatedness using *k*-mers, applying the same method as with SNPs, with presence/absence as the two alleles. We calculated the relatedness matrices for *A. thaliana*, maize, and tomato and compared them to the SNP-based relatedness. In all three species there was agreement between the two methods, although initial results were clearly better for *A. thaliana* and maize than for tomato (Fig. 6). The inferior performance in tomato was due to 21 accessions (Fig. S7), which appeared to be more distantly related to the other accessions based on *k*-mer than what had been estimated with SNPs. This is likely due to these accessions containing diverged genomic regions that do poorly in SNP mapping, resulting in inaccurate relatedness estimates. Removing these 21 accessions increased the correlation between SNP- and *k*-mer-based relatedness estimates from 0.60 to 0.83. In conclusion, *k*-mers can be used to calculate relatedness between individuals, thus paving the way for GWAS in organisms without high-quality reference genomes.

**Figure 6.**
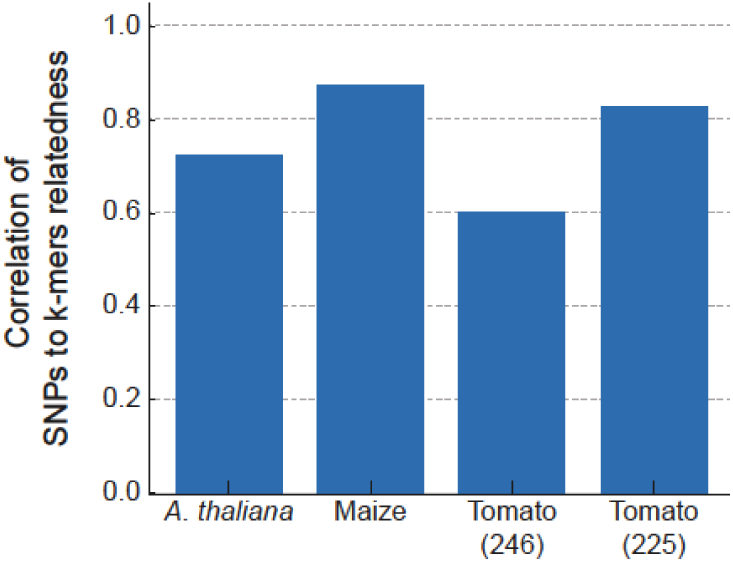
Kinship matrix estimates with *k*-mers. Relatedness between accessions was independently estimated based on SNPs and *k*-mers. The correlation between the two for tomato could be improved by removing 21 accessions that behaved differently between *k*-mers and SNPs (see Fig. S7).

## Discussion

The complexity of plant genomes makes identification of genotype-phenotype associations often challenging. To cope with this complexity, we followed a simple idea: most genetic variants leave a mark in the form of presence or absence of specific *k*-mers in whole genome sequencing data. Therefore, associating these *k*-mer marks with phenotypes will lead back to the genetic variants of interest. Our approach can identify associations found also by SNPs and short indels, but it excels when it comes to the detection of structural variants and variants not present in the reference genome. The expansion of variant types detected by our *k*-mer method complements SNP-based approaches, and greatly increases opportunities for finding and exploiting complex genetic variants driving phenotypic differences in plants, including improved genomic predictions.

*k*-mers mark genetic polymorphisms in the population, but the types and genomic positions of these polymorphisms are initially not known. While one can also use *k*-mers for predictive models without knowing their genomic context, in many cases the genomic contexts of *k*-mers associated with certain phenotypes are of interest. The simplest solution is to align the *k*-mers or the short reads they originate from to a reference genome, an approach that was effective for some phenotypes we have studied, as it has been in bacteria (Pascoe et al., 2015). However, if *k*-mers can be mapped to the reference genome, the underlying variants are likely to be also tagged by SNPs, as we saw for *A. thaliana* flowering time. In case *k*-mers cannot be placed on the reference genome, one can first identify the originating short reads and assemble these into larger fragments. We found this to be a very effective path to uncovering the genomic context of *k*-mers. Particularly the combination of reads from multiple accessions can provide high local coverage around the *k*-mers of interest, increasing the chances that sizeable fragments can be assembled and located in the reference genome or in other sequence databases. For example, in the GWA of days to tassel in maize, reads containing the associated *k*-mers could not be assigned to a specific location in the genome, but the assembled fragment mapped to a unique genomic position. This approach, manually applied in this study, provides a framework to systematically elucidate *k*-mer’s genomic context.

A main attraction of using *k*-mers as markers is that in principle they are able to tag many types of variants. A further improvement over our approach will be *k*-mers that tag heterozygous variants. In our current implementation, which relies on complete presence or absence of specific *k*-mers, only one of the homozygous states has to be clearly differentiated not only from the alternative homozygous state, but also from the heterozygous state. This did not affect comparisons between SNPs and *k*-mers in this study, as we only looked at inbred populations, where only homozygous, binary states are expected. Another improvement will be to use *k*-mers to detect causal copy number variations. So far, we can only tag copy number variants, if the junctions produce unique *k*-mers, but it would be desirable to use also *k*-mers inside copy number variants. Therefore, a future improvement will be an implementation that uses normalized counts instead of presence/absence of *k*-mers, which will create a framework that can, at least in principle, detect almost any kind of genomic variation.

The comparison of the *k*-mer- and SNP-based GWAS provides an interesting view on tradeoffs in the characterization of genetic variability. The stronger top p-values obtained with *k*-mers in cases where a SNP is the actual underlying genetic-variant points to incomplete use of existing information in SNP calling. On the other hand by minimizing filtering of *k*-mers, we included in our analysis some *k*-mers that represent only sequencing errors. Another potential source of noise comes from *k*-mers that are missed due to low coverage, which will be treated as absent. We reasoned that including these erroneous *k*-mers primarily has mostly computational costs, with some decrease in statistical power, since the chance of such *k*-mers generating an association signal is vanishingly small. Moreover, the high similarity of relatedness estimates using either SNPs (which are in essence largely filtered for sequencing errors) or all *k*-mers confirms that erroneous *k*-mers produce little signal. On the other hand, the higher effective number of *k*-mers compared to SNPs requires a more stringent threshold that takes the increased number of statistical tests into account and thereby decreases statistical power. This increase in test load is similar to the one that occurred when the genomics field moved from using microarray to next-generation sequencing in defining SNPs (1001 Genomes Consortium, 2016; The 1000 Genomes Project Consortium, 2010; Weigel and Mott, 2009). Thus, the higher threshold is an inevitable result from increasing our search space to catch more genetic variants.

*k*-mer associations inverts how GWAS is usually done. Instead of locating sequence variations in the genome and then associating them with a phenotype, we identify sequence-phenotype associations and only then find the genomic context of the sequence variations. Genome assemblies and genetic variant calling are procedures in which many logical decisions have to be made (Bradnam et al., 2013; Olson et al., 2015). These include high level decisions such as what information and software to use, as well as the many pragmatic thresholds chosen at each step of the way. Every community optimize these steps a bit differently, not least based on differences in the biology of the organisms they study, and surely these decisions affect downstream analyses (1001 Genomes Consortium, 2016; Bukowski et al., 2018; Tieman et al., 2017). Here, we took a complementary path in which initially neither a genome reference nor variant calling is needed, trying to reduce arbitrary decisions to a bare minimum. Technological improvement in short- and long-read sequences as well as methods to integrate them into a population-level genetic variation data-structure will expand the covered genetic variants (Paten et al., 2017; Schneeberger et al., 2009). While traditional GWAS methods will benefit from these technological improvements, so will *k*-mer based approaches, which will be able to use tags spanning larger genomic distances. Therefore, we posit that for GWAS purposes, *k*-mer based approaches are ideal because they minimize arbitrary choices when classifying alleles and because they capture more, almost optimal, information from raw sequencing data.

## Acknowledgment

We thank the many colleagues who have shared *A. thaliana* phenotypic information with us. We thank in particular G. Zhu and S. Huang for help with tomato genotypic and phenotypic information and C. Romay, R. Bukowski, and E. Buckler for help with maize genotypes and phenotypes. We thank K. Swarts, F. Rabanal, I Soifer, and R. Schweiger for fruitful discussions and comments on the manuscript. This work was supported by ERC AdG IMMUNEMENSIS, DFG ERA-CAPS “1001 Genomes Plus” and the Max Planck Society.

## Methods

### Curation of an *A. thaliana* phenotype compendium

Studies containing phenotypic data on *A. thaliana* accessions were located by searching NCBI PubMed using a set of general terms. For most studies, relevant data was obtained from the supplementary information or an online repository. Requests were sent to the corresponding authors of studies for which data could not be found in the public domain. Data already uploaded to the AraPheno dataset (Seren et al., 2017) downloaded from there. Phenotypic data in PDF format was extracted using Tabula software. Different sets of naming for accessions were converted to accession indices. In case an index for an accession could not be located, we omitted the corresponding data point. In case an accession could potentially be assigned to different indices, we first checked if it was part of the 1001 Genomes project; if so, we used the 1001 Genomes index. In case the accession was not part of it, one of the possible indices was assigned at random. Phenotypes of metabolite measurements from two studies, (Fordyce et al., 2018) and (Chan et al., 2010), were filtered to a reduced set by the following procedure: take the first phenotype, sequentially retain phenotypes if correlation with all previously taken phenotypes is lower than 0.7. Data from the second study (Chan et al., 2010), were further filtered for phenotypes with a title. Assignment of categories for each phenotype was done manually (Table S1). All processed phenotypic data can be found in Dataset S1.

### Whole genome sequencing data and variant calls of *A. thaliana*

Whole genome short reads for 1,135 *A. thaliana* accessions were downloaded from NCBI SRA (accession SRP056687). Accessions with fewer than 10^8^ unique *k*-mers, a proxy for low effective coverage, were removed, resulting in a set of 1,008 accessions. The 1001 Genomes project VCF file with SNPs and short indels was downloaded from http://1001genomes.org/data/GMI-MPI/releases/v3.1 and was condensed into these 1,008 accessions, using vcftools v0.1.15 (Danecek et al., 2011). We required a minimum minor allele count (MAC) of 5 individuals, resulting in 5,649,128 genetic variants. The VCF file was then converted to a PLINK binary file using PLINK v1.9 (Purcell et al., 2007). In case more than two alleles were possible in a specific location, PLINK keeps the reference allele and the most common alternative allele. The TAIR10 reference genome was used for short read and *k*-mer alignments. Coordinates for genes in figures were taken from Araport11 (Cheng et al., 2017).

### Whole genome sequencing data and variant calls of maize

Whole genome short reads of maize accessions corresponded to the “282 set” part of the maize HapMap3.2.1 project (Bukowski et al., 2018). Sequencing libraries “x2” and “x4” were downloaded from NCBI SRA (accession PRJNA389800) and combined. Coverage per accession was calculated as number of reads multiplied by read length and divided by the genome size, only data for 150 accessions with coverage >6x was used. Phenotypic data for 252 traits measured for these accessions were downloaded from Panzea (https://www.panzea.org) (Zhao et al., 2006).

Two of theses phenotypes were constant over more than 90% of the 150 accessions, these two were removed from further analysis (“NumberofTilleringPlants_env_07A”, “TilleringIndex-BorderPlant_env_07A”). The HapMap3.2.1 VCF files (c*_282_corrected_onHmp321.vcf.gz) of SNPs and indels were downloaded from Cyverse. Variant files were filtered using vcftools v0.1.15 to the relevant 150 accessions. Variants were further filtered for MAC of ≥5, resulting in a final set of 35,522,659 variants. The B73 reference genome, version AGPv3 (Portwood et al., 2019), that was used to create the VCF file was downloaded from MaizeGDB and used for short read and *k*-mer alignments(Portwood et al., 2019).

### Whole genome sequencing data and variant calls of tomato

Whole genome short reads were downloaded for 246 accessions with coverage >6x, from NCBI SRA and EBI ENA (accession numbers SRP045767, PRJEB5235 and PRJNA353161). A table with coverage per accession was shared by the authors (Tieman et al., 2017). Metabolite measurements were taken from (Tieman et al., 2017) (only adjusted values) and a subset of metabolites from (Zhu et al., 2018). These were filtered to a reduced set by the following procedure: take the first phenotype, sequentially retain phenotypes if correlation with all previously taken phenotypes is lower than 0.7. Metabolites were ordered as reported originally (Zhu et al., 2018). Only one repeat, the one with more data points and requiring at least 40 data points was retained. The VCF file with SNPs and short indels (Tieman et al., 2017) was obtained from the authors and filtered for the relevant 246 accessions. Variants were further filtered for MAC of ≥5, resulting in a final set of 2,076,690 variants. Reference genome SL2.5 (Tomato Genome Consortium, 2012) (https://www.ncbi.nlm.nih.gov/assembly/GCF_000188115.3/) used to create the VCF file was used for short read and *k*-mer alignments.

### *k*-mer counting and initial processing

For each accession from each of the three species all sequencing data from different runs were combined. The number of times each *k*-mer (k=25bp/31bp) appeared in the raw sequencing reads were counted using KMC v3 (Kokot et al., 2017). *k*-mers were counted twice, first counting canonical *k*-mers representation, which is the lower lexicographically for a *k*-mer and its reverse-complement. This list contains only *k*-mer appearing at least twice (maize and tomato) or thrice (*A. thaliana*) in the sequence reads. The second count includes all *k*-mers and without canonization. The KMC binary outputs of *k*-mers counts in the two lists were read using KMC C++ API, to keep all calculations in binary representation. For each *k*-mer in the first list, the information of which form (canonized, not-canonized, or both) it appeared in was extracted from the second list. This form information was coded in two bits, were the first/second bit indicates if the *k*-mer was observed in its canonized/non-canonized form, respectively. These two bits were inserted in the 2 most-significant-bits of the *k*-mer bit representation, as *k*-mers are of maximal length of 31bp, all information could be coded in a 64-bit word. The 64-bit *k*-mers representation were sorted according to the *k*-mer lexicographic order and saved to a file in binary representation.

For each species, the latter *k*-mers lists from all accessions were combined into one list according to the following criteria: only *k*-mers appearing in at least 5 accessions, and for a *k*-mer appearing in N accession it had to be observed in both canonized and non-canonized form in at least 0.2*N of the accessions. There were 2.26*10^9^, 2.21*10^9^, 3.23*10^9^, and 7.28*10^9^ unique *k*-mers in all accessions in the first type of counting, i.e. before filtering, and 439*10^6^, 393*10^6^, 981*10^6^, and 2.33*10^9^ passed the second criteria for *A. thaliana* (31-mers), *A. thaliana* (25-mers), tomato (31-mers), and maize (31-mers), respectively. The final filtered *k*-mers were outputted in binary format to a file, the histogram of number of *k*-mers appearances was calculated and saved during this process as well (e.g. Fig. S2A).

### Combining *k*-mers from different accessions to a *k*-mers presence/absence table

Tables containing the presence/absence per *k*-mer per accession in binary format were created, for each specie and *k*-mer size. The tables were organized as follows: *k*-mers information was written in serialized blocks of N+1 64-bit words. In each block, the first word codes the *k*-mer (k<32bp), the next N 64-bit blocks codes for the presence/absence of the *k*-mer in the different accessions: 1 in position i denoting the *k*-mer was found in accession i and 0 otherwise. N is the number of accessions divided by 64, rounded up. The last remaining padding bits not used were set to 0. Calculation of tables was done as follows: *k*-mers lists for all accession were opened together, in each step all the *k*-mers up to a threshold were read. *k*-mers were then combined in a sub-table to create the presence/absence patterns and then outputted in the described format with lexicographically ordered *k*-mers. This process was designed to minimize the memory load, and could be achieved due to the sorted *k*-mers in all separate lists.

### Counting and filtering unique presence absence patterns of *k*-mers

To check if a specific presence/absence pattern was already observed, the following method was used. This was done in order to count or filter the patterns. Each pattern, represented by a vector of N 64-bit words was inputted in a hash function which outputs a single 64-bit word. The hashed value was then stored in a set structure built on a hash-table. The size of the set was an indication of the number of unique patterns. Moreover, it was used continuously to filter patterns, by checking if a pattern (its hashed value) was already observed. The probability that two different patterns had the same hash value is very low: if we have *n* patterns, the space is of size *S* = 2^64^, the probability that at least one collision occurs randomly is:

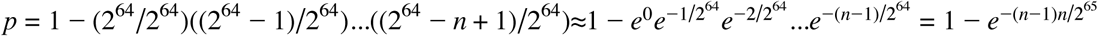

If *n* = 2^30^ > 1, 000, 000, 000 then 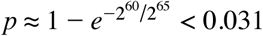, so there is ∼97% chance that not even one collision occurred for 1 billion distinct *k*-mers.

### Calculate and comparison of kinship matrices

Kinship matrix of relatedness between accessions was calculated as in EMMA (Kang et al., 2008), with default parameters. The algorithm was re-coded in C++ to read directly PLINK binary files for improved efficiency. For *k*-mers based relatedness the same algorithm was used, coding presence/absence as two alleles. For comparison of *k*-mers-to SNPs-based relatedness we correlated (pearson) the values for all 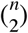 pairs, for *n* accessions. For tomato, 3492 pairs had a relatedness more than 0.15 lower for *k*-mer than for SNPs. 3,298 (94.4%) of these pairs were between a set of 21 accessions and all other 225 accessions. We calculated the correlation twice: for all pairs, and only between pairs of these 225 accessions.

### GWA on SNPs and short indels or on full *k*-mers table

Genome-wide association on the full set of SNPs and short indels was conducted using linear mixed models with the kinship matrix, using GEMMA version 0.96 (Zhou and Stephens, 2012). Minor allele frequency (MAF) was set to 5% and MAC was set to 5, with a maximum of 50% missing values (-miss 0.5). Kinship matrix was used to account for population structure. To run GWA on the full set of *k*-mers (e.g. in Fig. 1B), *k*-mers were first filtered for *k*-mers having only unique patterns on the relevant set of accessions, MAF of at least 5%, and MAC of at least 5. Presence/absence patterns were then condensed to only the relevant accessions and output as a PLINK binary file directly. GEMMA was then run using the same parameters as for the SNPs GWA described above.

### Phenotype covariance matrix estimation and phenotypes permutation

EMMA (emma.REMLE function) was used to calculate the variance components which were used to calculate the phenotypic covariance matrix (Kang et al., 2008). We then calculated 100 permutations of the phenotype using the mvnpermute R package (Abney, 2015). The n% (e.g. n=5 gives 5%) family-wise error rate threshold was defined by taking the n-th top p-value from the 100 top p-value of running GWA on each permutation. In all cases, unless indicated otherwise, where a threshold is referred to, it is the 5% threshold.

### Scoring p-values from GWA for similarity to uniform distribution and filtering phenotypes

Each SNP-based GWA run was scored for a general bias in p-value distribution, similar to Atwell et al. (Atwell et al., 2010). All SNPs p-values were collected, the 99% higher p-values were tested against the uniform distribution using a kolmogorov-smirnov test, and the test statistic was used to filter phenotypes for which distribution deviated significantly from the uniform distribution. A threshold of 0.05 was used, filtering 89, 0, and 295 phenotypes for *A. thaliana*, maize and tomato, respectively.

### *K*-mers genome-wide associations

Association of *k*-mers was done in two steps, with the aim of getting the most significant *k*-mers p-values. The first step was based on the approach used in Bolt-lmm-inf and GRAMMAR-Gamma (Loh et al., 2015; Svishcheva et al., 2012). For phenotypes *y*, genotypes *g*, and a covariance matrix Ω, the *k*-mer score is:

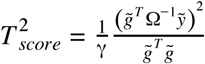

Where 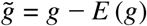 and 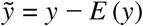. The first step was used only to filter a fixed number of top *k*-mers, thus we could use any score monotonous with 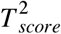, and specifically 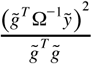 which is independent of γ (see supplementary note on calculation optimization). To keep used memory low, only best *k*-mers were stored in a priority queue data structure of constant size. The *k*-mers-table was uploaded to the memory in small chunks and associations were done with the phenotype and it’s permuted phenotypes for all *k*-mers in each chunk. The association step was implemented with the use of threads. After all *k*-mers were scored for associations with the phenotype and all its permutations, the *k*-mers-table was loaded again in chunks. The top *k*-mers with their genotype patterns were outputted in binary PLINK format, for the phenotype and each permutation separately. In the second step, the best *k*-mers were run using GEMMA to calculate the likelihood ratio test p-values (Zhou and Stephens, 2012).

The number of *k*-mers filter in the first step was set to 10,000 for *A. thaliana* and 100,000 for maize and tomato. Both steps associate *k*-mers while accounting for population structure, while the first step uses an approximation, the second use an exact model. Therefore, real top *k*-mers might be lost as they would not pass the first filtering step. To control for this, we first defined the 5% family-wise error-rate threshold based on the phenotype permutations, and then identified all the *k*-mers which passed the threshold. Next, we used the following criteria to minimize the chance of losing *k*-mers: we checked if all identified *k*-mers were in the top N/2 *k*-mers from the ordering of the first step (N=10,000 or 100,000 dependent on species). For example, in maize all *k*-mers passing the threshold in the second step should be in the top 50,000 *k*-mers from the first step. The probability that this will happen randomly is 2^−*m*^, where *m* is number of identified *k*-mers, in most phenotypes this is very unlikely. In 8.5% of phenotypes from *A. thaliana* the criteria was not fulfilled, for these phenotypes we re-run the two-steps with 100x more *k*-mers filtered in the first step, that is 1,000,000 *k*-mers. For 6 phenotypes the criteria still did not hold, these phenotypes were not used in further analysis. In tomato, 33% of phenotypes did not fulfill these criteria, in these cases we re-run the first step with 100x more *k*-mers filtered (10,000,000), 17 phenotypes still did not pass the threshold and were omitted from further analysis. The permutations were not re-run, and the threshold defined using 100,000 *k*-mers was used, as the top *k*-mer used to define the threshold tended to be high in the list. For maize all phenotypes passed the criteria and no re-running was needed.

### *Optimizing* of initial *k*-mers scoring

For: *N* – number of individuals, Ω – covariance matrix, *y* – phenotype, *g* – genotype (for *k*-mers taking the values 0 for absence and 1 for presence), and γ - GRAMMAR-Gamma factor which depends on the phenotype and relatedness between individuals, but not on specific *g* (Svishcheva et al., 2012).

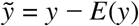 and 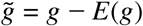

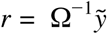 the transformed phenotype

The GRAMMAR-Gamma score of association 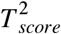 is distributed according to χ^2^ with 1 d.f. and satisfies:

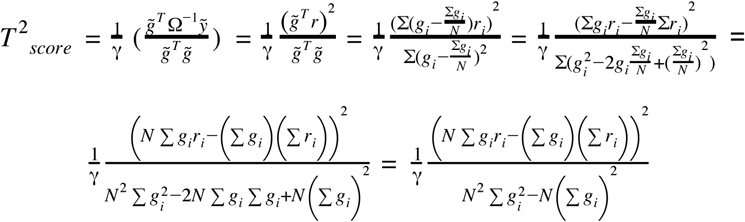

A *k*-mer can only be present or absent but not missing or heterozygous, thus 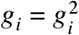 and we get:

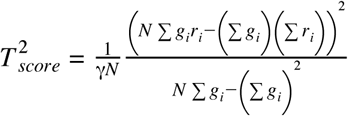

As we used the GRAMMAR-Gamma score only to filter the top *k*-mers, we did not need to calculate the p-value of 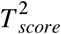 and could calculate a score that is monotonous with 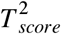, that is:

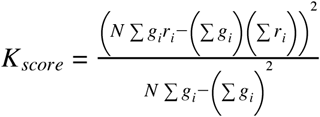

The summation ∑*r*_*i*_ can be calculated once per phenotype. Moreover, as we use permutation of phenotypes we can further optimize the scoring by calculating ∑*g*_*i*_ only once per *k*-mer.

For calculating the score of a specific *k*-mer, once ∑*r*_*i*_, ∑*g*_*i*_, and ∑*g*_*i*_*r*_*i*_ were calculated, we were left with 8 basic mathematical operations to obtain *K*_*score*_. Therefore, most of the computational load will be spent in the calculation of ∑*g*_*i*_*r*_*i*_, which requires 2*N* basic operations.

To computationally optimize the calculation of ∑*g*_*i*_*r*_*i*_, we used the Streaming SIMD Extensions 4 (SSE4) CPU instruction set. This implementation can be further optimized on a CPU that has AVX2, likely getting another 2-fold increase in efficiency with only small modifications to the code, however, we have not tested this option.

To optimize the GRAMMAR-Gamma filtering of SNPs we cannot benefit from the same optimizations as for *k*-mers. This is due to missing and heterozygous values a SNP can take. Therefore, in this case 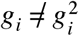. For SNPs our score will take the same form as 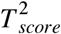:

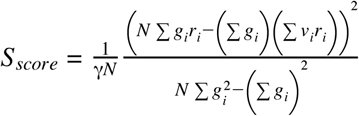

In this case *N* is different for different SNPs, and so as ∑*r*_*i*_. This later summation can be written as ∑*v*_*i*_*r*_*i*_, by defining *v*_*i*_ = 0 for *gi* = *missing* and *v*_*i*_ = 1 for *g*_*i*_ ≠ *missing*.

Thus, ∑*v*_*i*_*r*_*i*_ is specific for each SNP’s score and as *g*_*i*_ can also get the value 0.5, we separated ∑*g*_*i*_*r*_*i*_ to two separate dot-products in our implementation, as genotypes are coded by bit vectors.

### SNPs-based GWAS on phenotype permutations

To calculate thresholds for SNPs-based GWAS we used the two step approach used for *k*-mers. The permuted phenotypes were run in two steps as we were only interested in the top p-value to define thresholds. We filtered 10,000 variants in the first step which were then run using GEMMA to get exact scores (Zhou and Stephens, 2012). The non-permuted phenotype were run using GEMMA on all the variants.

### Calculation of linkage-disequilibrium (LD)

For two variants, *x* and *y*, each can be a *k*-mer or a SNP, LD measure was calculated using the r^2^ measure (Devlin and Risch, 1995). For a *k*-mer, variants were coded as 0/1, if absent or present, respectively. For SNPs one variant was coded as 0 and the other as 1. If one of the variants had a missing or heterozygous value in a position, this position was not used in the analysis. The LD value was calculated using the formula:

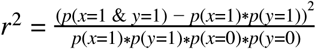

### LD cumulative graph (Fig 2E,H)

For a set of phenotypes and for every *l* = 0, 0.05,.., 1 we calculated the percentage of phenotypes for which exists a *k*-mer or a SNP in the pre-defined group which is in *LD*≥*l* with top SNP or top *k*-mer, respectively. The pre-defined groups are: (1) all the *k*-mers which passed the SNPs defined threshold in Figure 2E or (2) all the SNPs or *k*-mers which passed their own defined thresholds in Figure 2H. The percentage is then plotted as a function of *l*.

### Retrieving source reads of a specific *k*-mer and assembling them

For a *k*-mer identified as being associated with a phenotype we first looked in the *k*-mers-table and identified all accessions taking part in the association analysis and having this *k*-mer present. For each of these accessions we went over all sequencing reads and filtered out all paired-end reads which contained the *k*-mer or its reverse-complement. To assemble paired-reads, SPAdes v3.11.1 was used with “--careful” parameter (Bankevich et al., 2012).

### Alignment of reads or *k*-mers to the genome

Paired-end reads were aligned to the genome using bowtie2 v2.2.3, with the “--very-sensitive-local” parameter. *k*-mers were aligned to the genome using bowtie v1.2.2 with “--best --all --strata” parameters (Langmead and Salzberg, 2012).

### Analysis of flowering time in 10C (Figure 1, Figure S2)

To find the location in the genome of the 105 identified *k*-mers, *k*-mers were first mapped to the *A. thaliana* genome. 84 of the *k*-mers had a unique mapping, one *k*-mer was mapped to multiple locations and 20 could not be mapped. For the 21 *k*-mers with no unique mapping we located the sequencing reads they originated from, and mapped the reads to the *A. thaliana* genome. For each of the *k*-mers we looked only on the reads with the top mapping scores. For the one *k*-mer which had multiple possible alignment also the originating reads did not have a consensus mapping location in the genome. For every *k*-mer from the 20 non-mapped *k*-mers, all top reads per *k*-mer, in some cases except one, mapped to a specific region spanning a few hundred base pairs. The middle of this region was defined as the *k*-mer position for the Manhattan plot in Figure 1D.

To find the location of the 93 associated *k*-mers of length 25bp, presented in supplementary Figure S2D, we followed the same procedure. 87 of the *k*-mers had a unique mapping, one was mapped multiple times and 5 could not be mapped. For the 5 *k*-mers with no mapping and the *k*-mer with non unique mapping, we located the originated short reads and aligned them to the genome. For each of the 5 *k*-mers with no mapping, all reads with top mapping score mapped to a specific region of a few hundred base pairs, we took the middle of the region as the *k*-mer location in the Manhattan plot. For the *k*-mer with multiple mappings, 15 out of the 17 reads mapped to the same region and we used this location. All *k*-mers mapped to the 4 location in the genome for which SNPs were identified except one - AAGCTACTTGGTTGATAATACTAAT. The reads from which this *k*-mer originated mapped to the same region in chromosome 5 position 3191745-3192193 and we used the middle of this region as the *k*-mer location.

### Analysis of xylosides percentage (Figure 3A,B)

All *k*-mers passing the threshold, were mapped uniquely to chromosome 5 in the region 871,976 – 886,983. Of the 123 identified *k*-mers, 27 had the same minimal p-value (*log*10 (*p value*) = 44.7). These *k*-mers mapped to chromosome 5 in positions 871,976 to 872,002, all covering the region 872,002-872,007. For the 60 accessions used in this analysis, all reads from the 1001G were mapped to the reference genome. The mapping in region 872,002-872,007 of chromosome 5 were examined manually by IGV in all accessions (Robinson et al., 2011), and the 2 SNPs 872,003 and 872,007 were called manually without knowledge of the phenotype value.

### Analysis of growth inhibition in presence of flg22 (Figure 3C,D)

The phenotype in the original study was labeled “flgPsHRp” (Vetter et al., 2016). For each of the 7 *k*-mers which could not be mapped uniquely to the genome, the originated reads from all accessions were retrieved and assembled. All the seven cases resulted in the same assembled fragment (SEQ1, table S2). Using NCBI BLAST we mapped this fragment to chromosome 1: position 40-265 were mapped to 8169229-8169455 and position 262-604 were mapped to 8170348-8170687. For every accession from the 106 that were used in the GWAS analysis we tried to locally assemble this region, to see if the junction between chromosome 1 8169455 to 8170348 could be identified. We used all the 31bp *k*-mers from the above assembled fragment as bait, and located all the reads for each accession separately. For 11 out of the 13 accessions that had all 10 identified *k*-mers we got a fragment from the assembly process. In all 11 cases the exact same junction was identified. For 1 of the 4 accessions that had only part of the 10 identified *k*-mer we got a fragment from the assembler, which had the same junction. For 43 of the 89 accessions that had none of the identified *k*-mers the assembly process resulted in a fragment, in none of these cases the above junction could be identified.

### Analysis of germination in darkness and low nutrients (Figure 3E, F)

The phenotype in the original study was labeled “k_light_0_nutrient_0” (Morrison and Linder, 2014). The 11 identified *k*-mers had two possible presence/absence patterns, separating them into two groups of 4 and 7 *k*-mers. The short-read sequences containing the 4 or 7 *k*-mers were collected separately and assembled, resulting in the same 458bp fragment (SEQ2, table S2). This fragment was used as a query in NCBI BLAST search, resulting in alignment to Ler-0 chromosome 3 (LR215054.1) positions 15969670 to 15970128. The LR215054.1 sequence was downloaded and the region between (15969670-3000) to (15970128+3000) was retrieved and used as query to a NCBI BLAST search. The BLAST search resulted in a mapping to Col-0 reference genome chromosome 3 (CP002686.1). Region 1-604 mapped to 16075369-16075968, region 930-1445 mapped to 16076025-16076532, region 3446-3946 mapped to 16079744-16080244, and region 3958-6459 mapped to 16080301-16082781.

### Analysis of root branching zone (Figure 3G)

The phenotype in the original study was labeled “Mean(R)_C”, that is Branching zone in no treatment (Ristova et al., 2018). No SNPs and 1 *k*-mer (AGCTACTTTGCCACCCACTGCTACTAACTCG) passed their corresponding 5% thresholds. The *k*-mer mapped the chloroplast genome in position 40297, with 1 mismatch. No SNPs and another *k*-mer (CCGGCGATTACTAGAGATTCCGGCTTCATGC) passed the 10% family-wise error-rate threshold. This *k*-mer mapped non-uniquely to two place in the chloroplast genome: 102285 and 136332.

### Analysis of Lesion by *Botrytis cinerea* UKRazz (Figure S3A)

The Lesion by *Botrytis cinerea* UKRazz phenotype was labeled as “Lesion_redgrn_m_theta_UKRazz”. In the GWAS analysis 19 *k*-mers and no SNPs were identified. All *k*-mers had the same presence/absence pattern. The short-read sequences from which the *k*-mers originated were mapped to chromosome 3 around position 72,000bp, and contained a 1-bp deletion of a T nucleotide in position 72,017. Whole genome sequencing reads were mapped to the genome for the 61 accessions with phenotypes used in these analyses. We manually observed the alignment around position 72,017 of chromosome 3, without the prior knowledge if the accession had the identified *k*-mers. For 20 accessions, we observed the 1-bp deletion in position 72,017, all 19 accessions containing the *k*-mers were part of these 20.

### Analysis of days to tassel and ear weight in maize (Figure 4)

Ear weight phenotype was labeled “EarWeight_env_07A” in original dataset (Zhao et al., 2006). Days to tassel was measured in growing degree days (GDD) and was labeled as “GDDDaystoTassel_env_06FL1” in original dataset. In comparison of LD between *k*-mers and SNPs in days to tassel (Fig. 4E, upper panel), two SNPs were filtered out as having more than 10% heterozygosity and one as having, exactly, 50% missing values. In days to tassel the *k*-mer which was similar to identified SNPs was AGAAGATATCTTATGAACTCCTCACCAGTAA. The 171 paired-end reads from which this *k*-mer originated mapped to the genome as follows - 2 (1.17%) aligned concordantly 0 times, 2 (1.17%) aligned concordantly exactly 1 time, and 167 (97.66%) aligned concordantly >1 times. The assembly of these reads produce two fragments, the first of length 273bp with coverage of 1.23 and the second of length 924bp and with coverage of 27.41 (SEQ3, table S2). We aligned this fragment to the genome using Minimap2, with the default parameters (Li, 2018). Minimap2 reported only 1 hit to chromosome 3 (NC_024461.1) in positions 159141222-159142137.

### Analysis of guaiacol concentration in tomato (Figure 5)

Guaiacol concentration was labeled “log3_guaiacol” in the original study. From the 293 *k*-mers passing the threshold, 184 could be mapped uniquely to the genome: 135 to chromosome 0 between position 12573795-12576534, and 45 to chromosome 9 between position 69301436-69305717, 3 to chromosome 6 between position 8476136-8476138, and 1 to chromosome 4 at position 53222324. The 4 *k*-mers mapped to chromosome 4 and 6 were checked manually by locating the reads containing them and aligning the reads to the genome, in all cases no reads were able to be aligned to the genome (>99.5% of reads). For the 35 *k*-mers not mapping to genome and in high LD, visualized in Figure 5E, all reads containing at least one of the *k*-mers were retrieved and assembled (SEQ4, table S2). NCBI Blast search of this fragment resulted in: positions 1-574 mapped to positions: 12578806-12579379 in chromosome 0 of the tomato genome (CP023756.1) and positions 580-1169 mapped to positions 289-878 in NSGT1 (KC696865.1). The R104 “smoky” accession NSGT1 ORF starts at position 307, as reported previously (Tikunov et al., 2013). NCBI BLAST of NSGT1 (KC696865.1), identified mapping to chromosome 9 of the tomato genome (CP023765.1), from positions 975-1353 to positions 69310153-69309775.

## Code availability

Code is available in https://github.com/voichek/kmersGWAS.

## Supplementary materials

**Figure S1:**
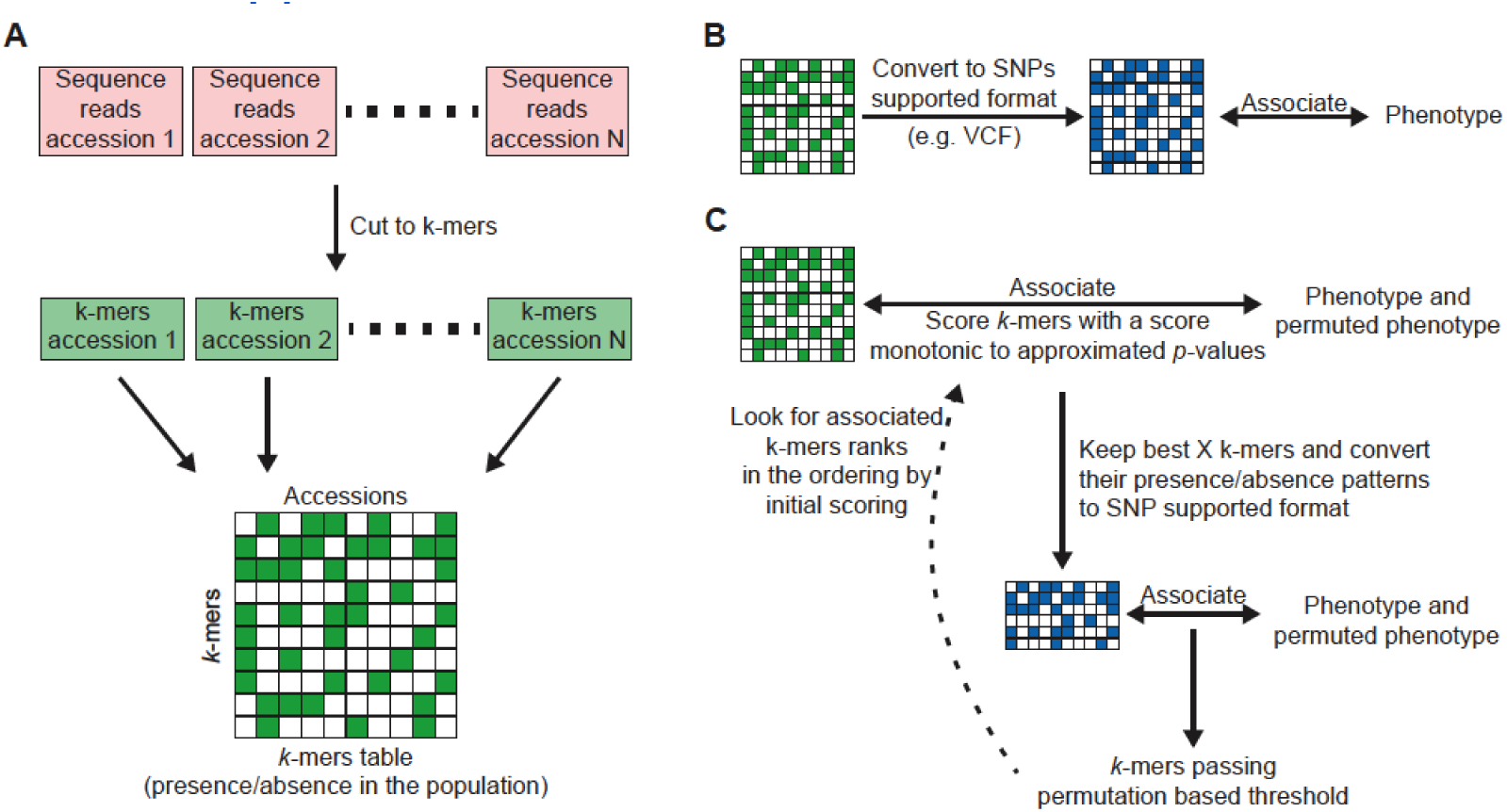
Scheme of pipeline for *k*-mer-based GWAS. **(A)** Creating the *k*-mer presence/absence table: Each accession’s genomic DNA sequencing reads are cut into *k*-mers of constant length using KMC (Kokot et al., 2017). Only *k*-mers appearing at least twice/thrice in a sequencing library are used. *k*-mers are further filtered to retain only those present in at least 5 accessions, and ones that are also found in their reverse-complement form in at least 20% of accessions they appear in. *k*-mer lists from all accessions are then combined into a *k*-mer presence/absence table. This table is encoded in a binary format, with each cell represented as a single bit. **(B)** Genome-wide associations on the full *k*-mer table using SNP-based software the: *k*-mer table can be converted into PLINK binary format, which can be used directly as input for association mapping in various software for SNP-based GWA (Purcell et al., 2007; Zhou and Stephens, 2012). **(C)** GWA optimized for the *k*-mers presence/absence table: *k*-mers presence/absence patterns are first associated with the phenotype and its permutations using a linear-mixed model to account for population structure (Loh et al., 2015; Svishcheva et al., 2012). This first step is done by calculating a score monotonic to an approximation of the exact model. This scoring system is ultra-fast and is built for the high computational load coming from the large number of *k*-mers and many permutations of phenotypes. Best *k*-mers from this first step (e.g. 100,000 *k*-mers) are used in the second step. In the second step an exact *p*-value is calculated (Zhou and Stephens, 2012) for all *k*-mers for both the phenotype and its permutations. A permutation-based threshold is calculated and all *k*-mers passing this threshold are checked for their rank in the scoring from the first step. If not all *k*-mers hits are in the top 50% of the initial scoring, then the entire process is rerun from the beginning, passing more *k*-mers from the first to the second step. This last test is built to confirm that the approximation of the first step will not remove true associated *k*-mers.

**Figure S2:**
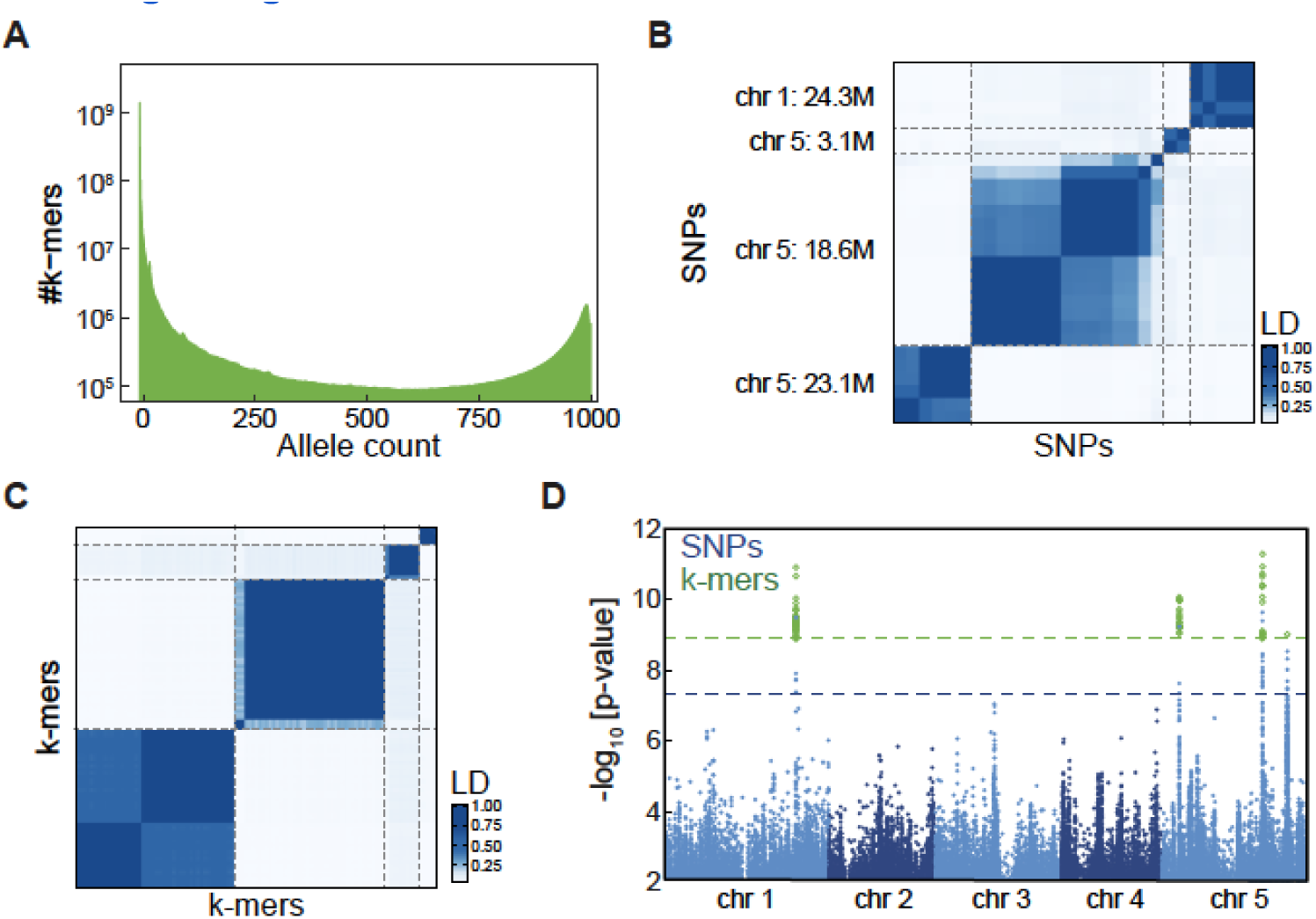
Flowering time genetic associations in *A. thaliana* identified with *k*-mers. **(A)** Histogram of *k*-mer allele counts: For every N=1..1008, plotted how many *k*-mers appeared in exactly N accessions. **(B)** LD between SNPs associated with flowering time. Dashed lines represent the four variant types, as in Figure 1C. **(C)** LD between *k*-mers associated with flowering time, Dashed lines represent the four variant types, as in Figure 1C. **(D)** Manhattan plot of SNPs and *k*-mer associations with flowering time in 10°C as in Figure 1D for *k*-mers of length 25bp.

**Figure S3:**
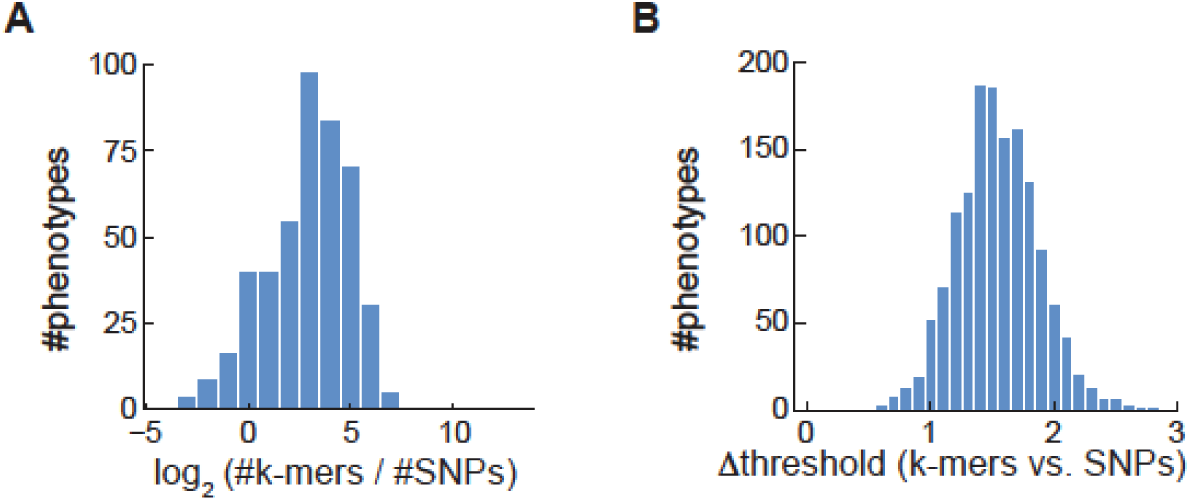
Comparison of SNP- and *k*-mer-GWAS on phenotypes from 104 studies on *A. thaliana* accessions. **(A)** Histogram of the number of identified *k*-mers vs. identified SNPs (in log_2_) for *A. thaliana* phenotypes. Only the 458 phenotypes with both variant types identified were used. **(B)** Histogram of thresholds difference of *k*-mers vs. SNPs of all *A. thaliana* phenotypes. Thresholds were -log_10_ transformed.

**Figure S4:**
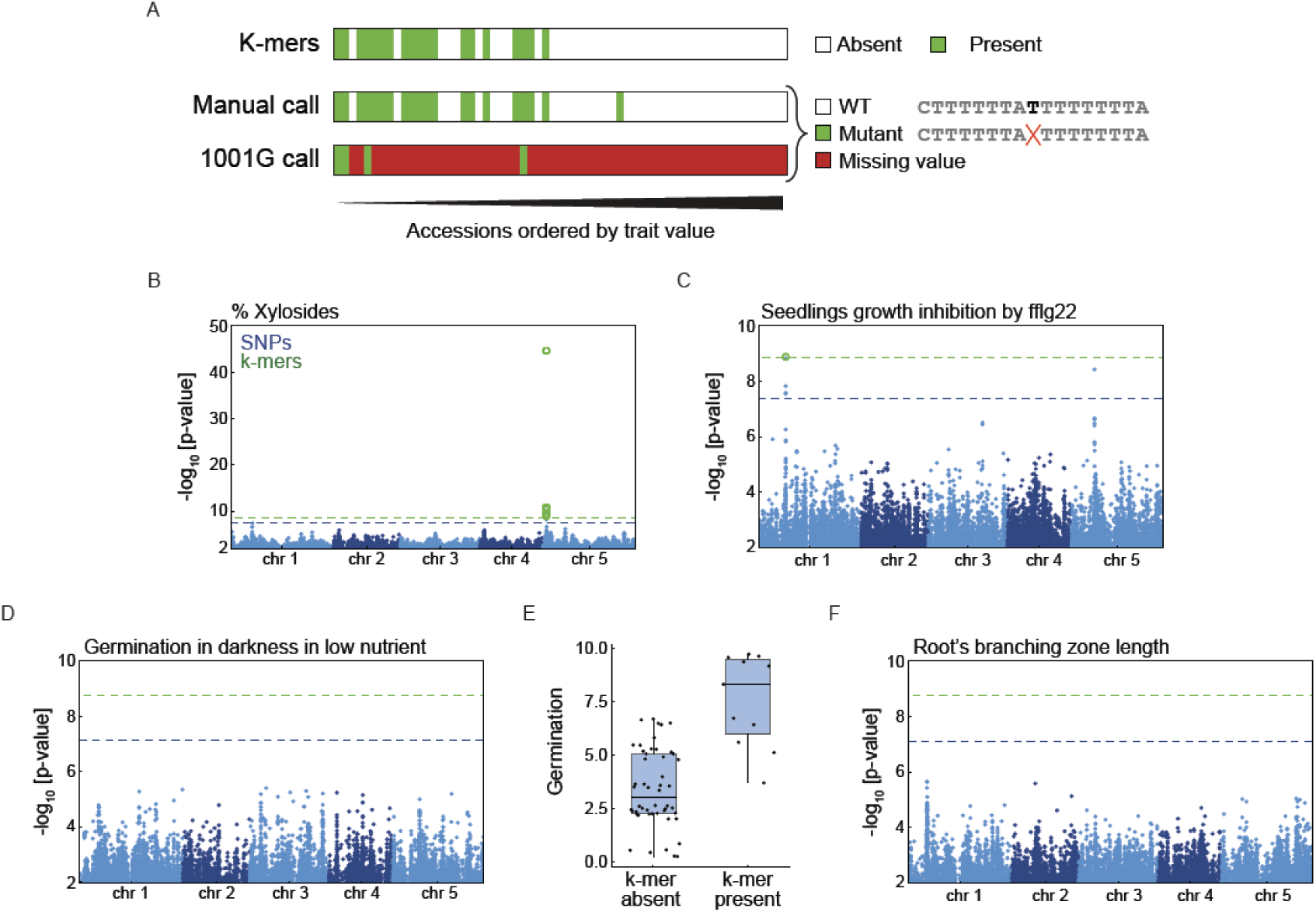
Specific case studies in which *k*-mers are superior to SNPs. **(A)** Results from GWAS on measurements of lesion by *Botrytis cinerea* UKRazz strain (Fordyce et al., 2018), an example of *k*-mers having better hold on genetic-variants present in the SNPs/indels table. We identified 19 *k*-mers and no SNPs as being associated with this phenotype. All the *k*-mers had the same presence/absence pattern (top row). The short sequence reads containing the *k*-mers mapped to chromosome 3 in proximity to position 72,000. The reads contained a single T nucleotide deletion in position 72,017, relative to the reference genome. The T nucleotide was part of an 8 T’s strach, the reference and mutated sequence around the deletion are indicated to the right of the manual calling for all accessions (middle row) and to the calls from the 1001G project (bottom). In the 1001G only 4 accessions were called out of the 61 accessions part of the analysis, for the other accessions, the tabled contained missing values. **(B)** Manhattan plot, for xyloside percentage. A focused view on region with identified associations is presented in Figure 3A. **(C)** Manhattan plot, for seedling growth inhibition by flg22. A focused view on region with identified associations is presented in Figure 3C. **(D)** Manhattan plot, for germination in darkness in low nutrient conditions. All identified *k*-mers could not be mapped to the genome. **(E)** The germination phenotype is plotted for accessions which have the top associated *k*-mer and those that do not. **(F)** Manhattan plot, for root’s branching zone length. Identified *k*-mer mapped the chloroplast genome, and thus not present in the graph.

**Figure S5:**
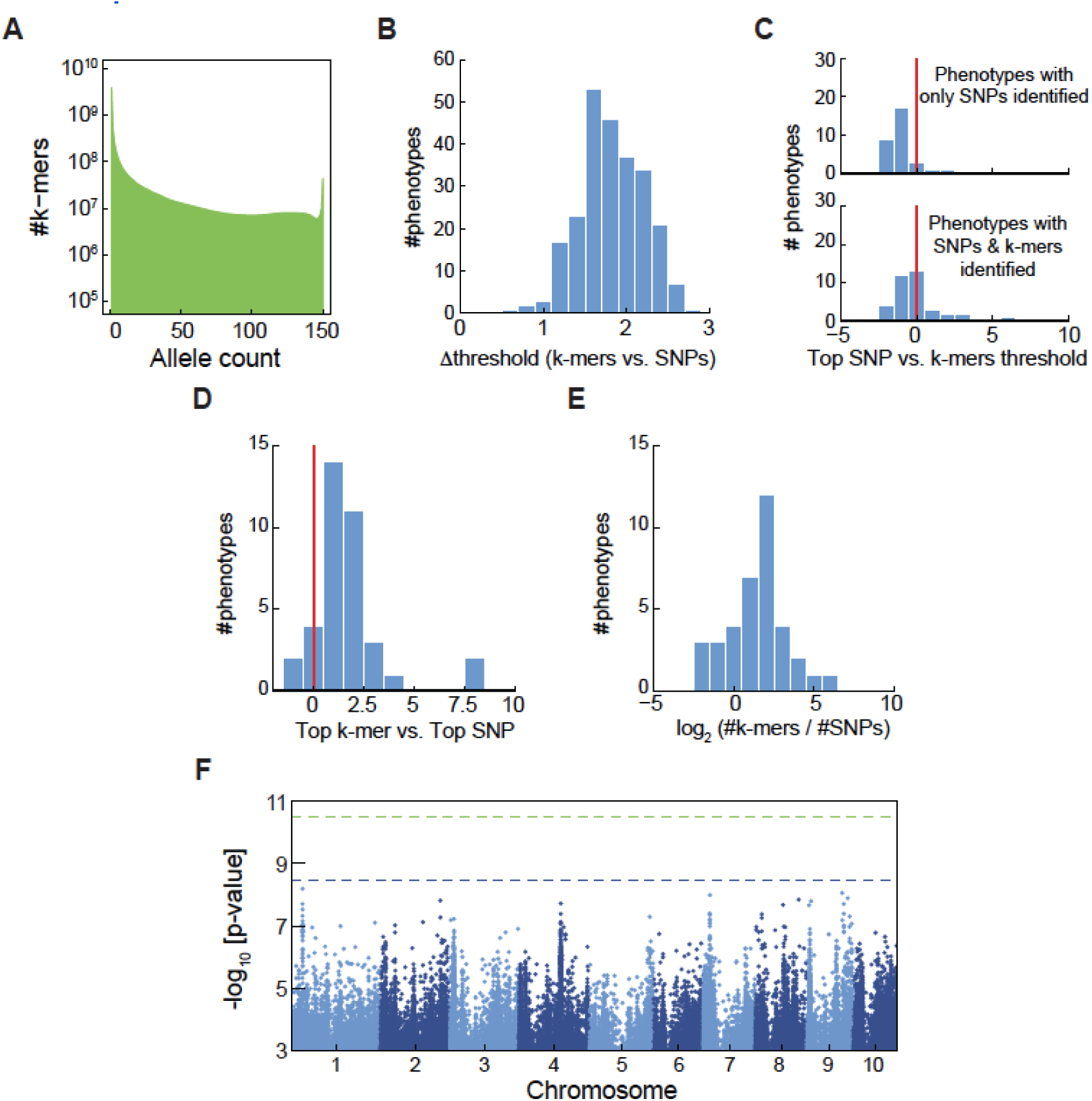
Comparison of SNP- and *k*-mer-based GWAS in maize. **(A)** Histogram of *k*-mer allele counts for maize accessions. **(B)** Histogram of difference between threshold values of SNPs and *k*-mers for maize phenotypes. **(C)** Histogram of the top SNP p-value divided by the *k*-mers defined threshold, in (-log10), for maize phenotypes. Plotted for phenotypes with only identified SNPs (upper panel) or for phenotypes with both SNPs and *k*-mers identified (lower panel). **(D)** Histogram of the difference between top (-log10) p-values in the two methods for maize phenotypes identified by both methods. Plottes as in Figure 2G. **(E)** Histogram of the number of identified *k*-mers vs. identified SNPs for maize phenotypes. **(F)** Manhattan plot of association with ear weight (environment 07A). Associated *k*-mers genomic location were not located, and are thus not presented.

**Figure S6:**
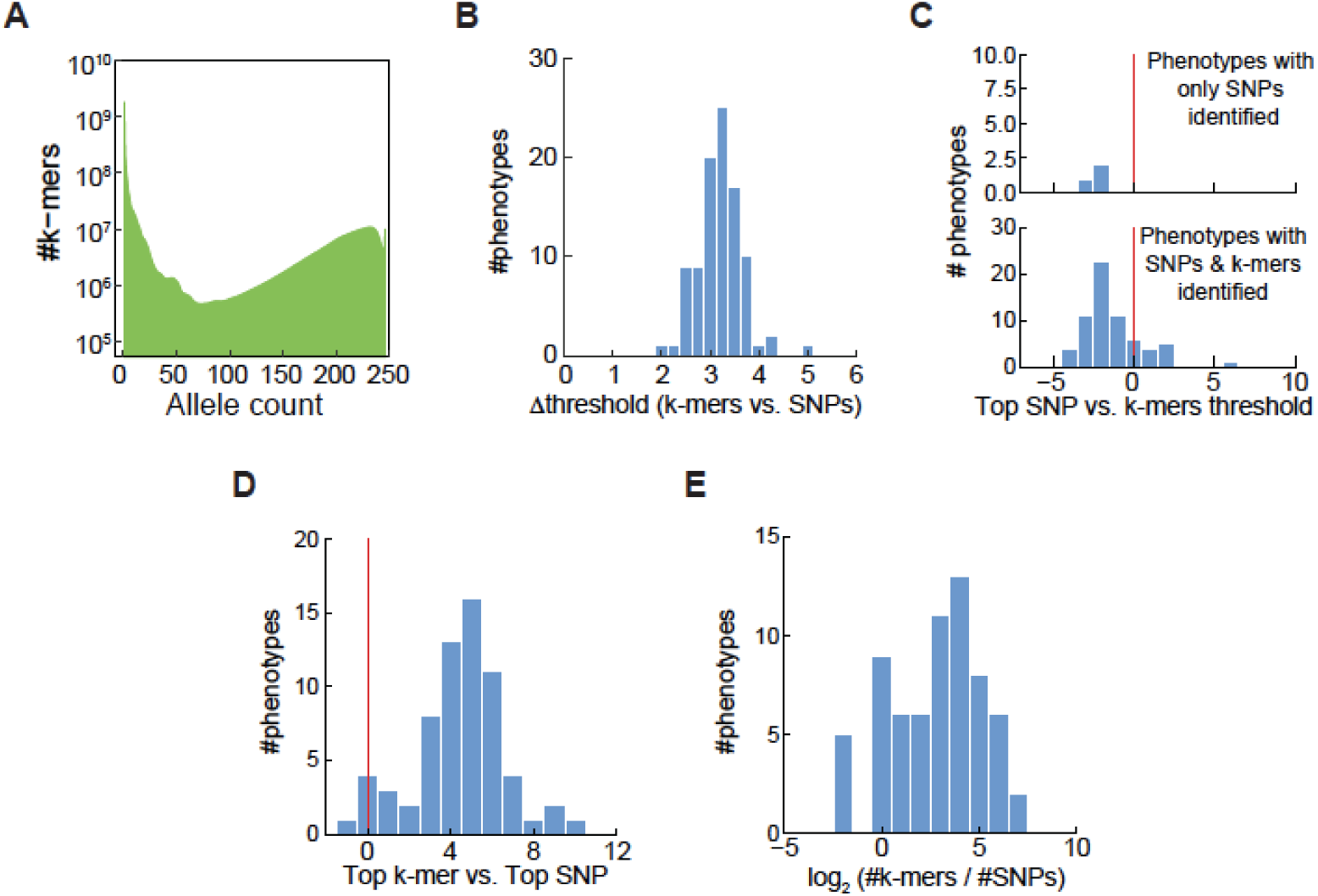
Comparison of SNP- and *k*-mer-based GWAS in tomato. **(A)** Histogram of *k*-mers allele counts for tomato accessions. **(B)** Histogram of difference between threshold values of SNPs and *k*-mers for tomato phenotypes. **(C)** Histogram of the top SNP p-value divided by the *k*-mers defined threshold, in (-log10), for tomato phenotypes. Plotted for phenotypes with only identified SNPs (upper panel) or for phenotypes with both SNPs and *k*-mers identified (lower panel). **(D)** Histogram of the difference between top (-log10) p-values in the two methods for tomato phenotypes. **(E)** Histogram of the number of identified *k*-mers vs. identified SNPs for tomato phenotypes.

**Figure S7:**
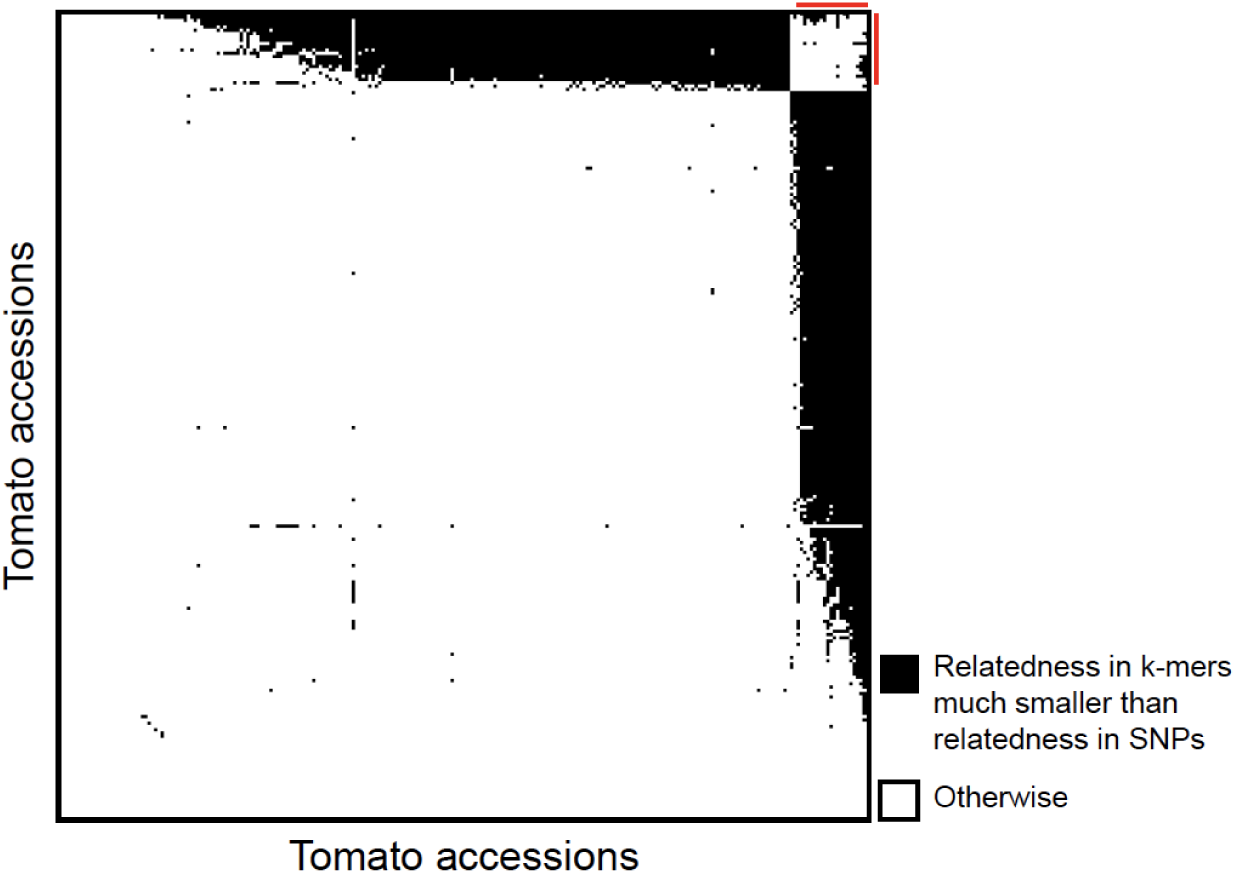
Kinship matrix calculation based on *k*-mers. Identification of pairs of tomato accessions for which relatedness as measured with *k*-mers is much lower than relatedness as measured with SNPs. For every pair among the 246 accessions, a black square is plotted if the difference in relatedness between SNPs and *k*-mers is larger than 0.15. Accessions are ordered by the number of black square in their row/column. Red lines mark the 21 accessions with most black squares, that is, those for which the *k*-mer/SNP difference in relatedness is larger than 0.15 for the most pairs.

**Table S2:**
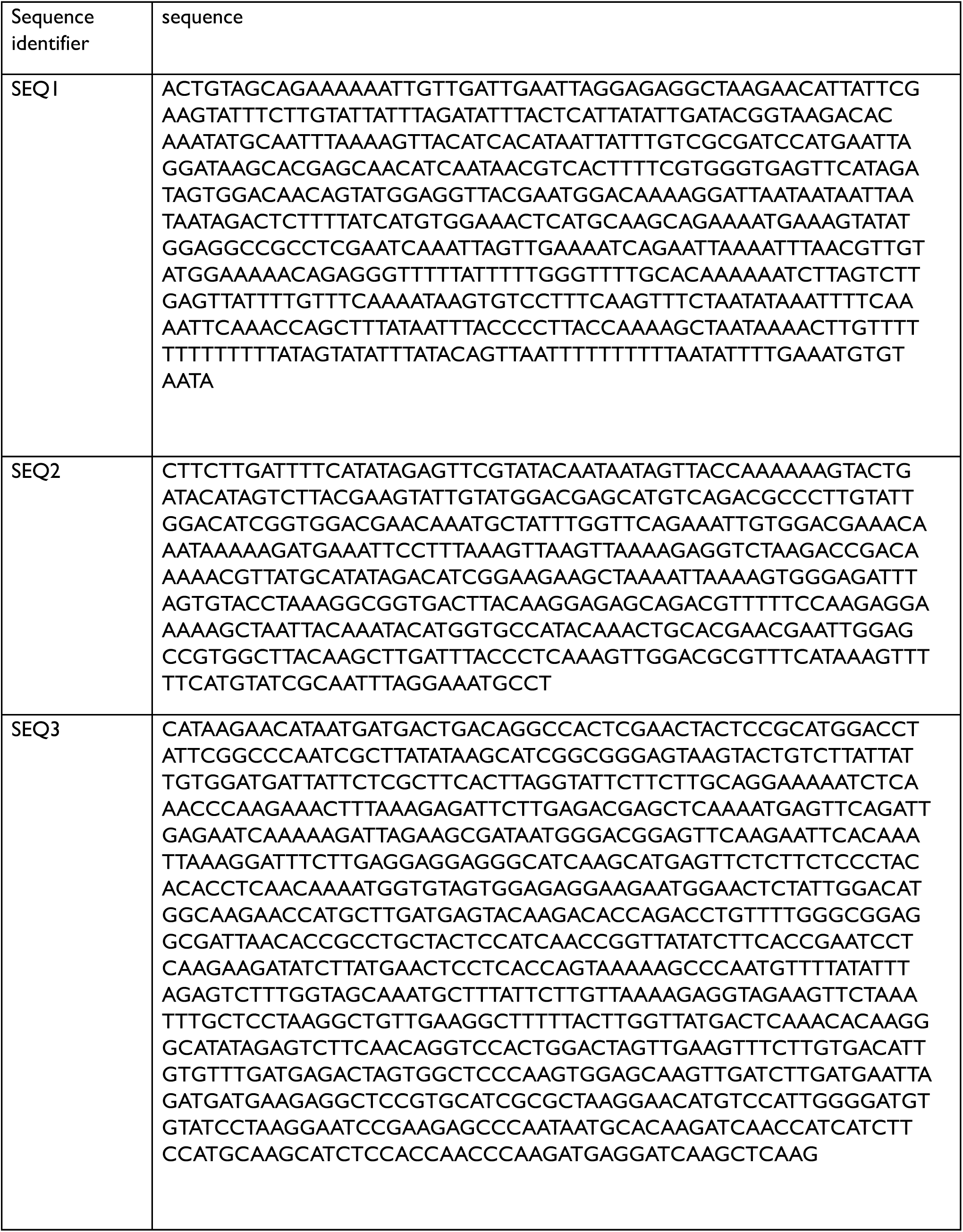
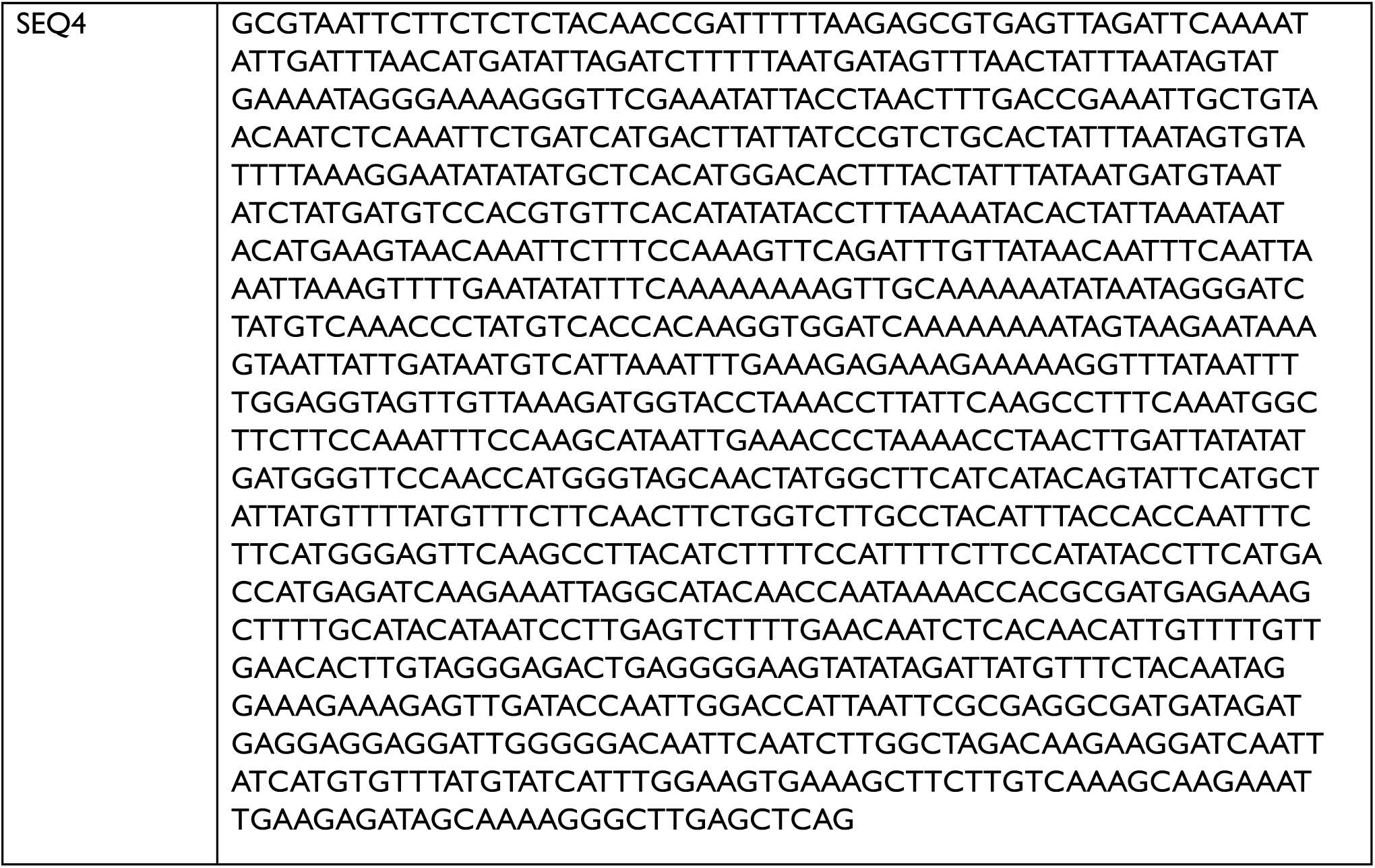
Assembled fragments from retrieved reads.

