## Supplementary material for "Finding genetic variants in plants without complete genomes": Dataset S1: README

Supplementary dataset 1 - Collection of phenotypes and top SNPs/k-mers passing their threshold.

Each phenotype has a separate directory which contains:
1) original_phenotype_values.txt - the phenotypic values for each accession. For some of the A. thaliana’s phenotypes, the accession in the original work were marked not by their index, but with some other identifier. In these cases we had to find the accession’s index as explained in the paper’s method. For the cases we could not find the index, the phenotypic value was not used, and will be missing from this file. 
2) used_phenotype_values.txt - a subset of the original_phenotype_values.txt file. In this file we only include accessions for which we have sequencing data for. This are the accessions we used for the GWAS analysis.

If the phenotype was not filtered out, the directory will contain additional 4 files:
3) kmers_pass_5percent_threshold_GEMMA_results.txt - The association results, as calculated by GEMMA, for all k-mers passing the 5% threshold.
4) kmers_pass_10percent_threshold_GEMMA_results.txt - The association results, as calculated by GEMMA, for all k-mers passing the 10% threshold.
5) snps_pass_5percent_threshold_GEMMA_results.txt - The association results, as calculated by GEMMA, for all SNPs passing the 5% threshold.
6) snps_pass_10percent_threshold_GEMMA_results.txt - The association results, as calculated by GEMMA, for all SNPs passing the 10% threshold.
